# Temporal–orbitofrontal pathway regulates choices across physical reward and visual novelty

**DOI:** 10.1101/2025.04.15.648986

**Authors:** Takaya Ogasawara, Kevin Xu, Abraham Z. Snyder, Joel S. Perlmutter, Zhude Tu, Takafumi Minamimoto, Ken-ichi Inoue, Masahiko Takada, Ilya E. Monosov

## Abstract

Perceptually novel objects have profound impacts on our daily decisions. People often pay to try novel meals over familiar ones, or to see novel visual scenes at art exhibits and travel destinations. This suggests that perceptual novelty and the value of physical rewards, such as food, interact at the level of neural circuits to guide decisions, but where and how is unknown. We designed a behavioral task to study this novelty-reward interaction in animals and to uncover its neural underpinnings. Subjects chose among familiar offers associated with different expectations of novel objects and different expectations of juice rewards. Expectation of novel objects increased the preference for expected rewards. This novelty-reward interaction was reflected by neural activity in the anterior ventral temporal cortex (AVMTC) – a region previously implicated in the detection and prediction of novelty – and in the orbitofrontal cortex (OFC) – an area that receives prominent AVMTC inputs and is known for its capacity to signal subjective value of visual objects. Neural activity patterns suggested that AVMTC was upstream of OFC in the decision process. Chemogenetic disruption of the AVMTC**→**OFC circuit altered the impact of expected novelty on the valuation of physical reward. Hence, the ventral visual system impacts novelty-reward interactions during decisions through direct projections to OFC.

## INTRODUCTION

Perceptually novel objects profoundly impact our daily decision-making. For example, commercial advertisements promote new products, such as clothes, cosmetics, and cell phones, and many people choose to purchase novel products over proven familiar ones for novelty’s own sake, sometimes over-paying for material goods and bearing additional costs. However, despite the ubiquity of these behaviors and their importance in our daily lives, how novelty interacts with reward remains poorly understood, particularly at the level of neural circuits. This lack of knowledge impacts our ability to develop a mechanistic understanding of psychiatric disorders in which novelty-seeking often goes awry, producing maladaptive behavioral and internal states (Djamshidian et al., 2011; Donfrancesco et al., 2015; S. W. Kim & Grant, 2001; Kusunoki et al., 2000; Young et al., 1995).

Theories regarding how novelty could impact the valuation of physical reward come from emerging developments in machine learning, largely inspired by psychology, economics, and education research (Aubret et al., 2023; Berlyne, 1950, 1954, 1966; Caplin & Dean, 2015; Caplin & Leahy, 2001; Cohen et al., 2007; Dubey & Griffiths, 2020; Gittins et al., 2011; Gottlieb et al., 2016; Gottlieb & Oudeyer, 2018; Hall & Smith, 1903; Jaegle et al., 2019; James, 1913, 1983; Kahneman & Tversky, 1979; Knight, 1921; Kreitler et al., 1975; Kreps & Porteus, 1978; Modirshanechi, Kondrakiewicz, et al., 2023; Murayama et al., 2019; Poli et al., 2024; Smock & Holt, 1962). For example, artificial agents that endow novelty with value have a behavioral advantage in many foraging environments (Aubret et al., 2023; Bellemare et al., 2016; Burda et al., 2018; Gottlieb et al., 2016; Houillon et al., 2013; Jaegle et al., 2019; Kakade & Dayan, 2002; Y. Kim et al., 2019; Modirshanechi, Kondrakiewicz, et al., 2023; Modirshanechi, Lin, et al., 2023; Oudeyer et al., 2007; Poli et al., 2024; Savinov et al., 2019; Tang et al., 2017; H. A. Xu et al., 2021). However, biologically inspired solutions to the question of *how* to endow novelty with value remain elusive. For example, we do not yet know if and how novelty impacts the valuation of physical reward at the level of neuronal activity, that is, whether it directly modulates the neuronal representation of physical reward value during choice.

To illuminate this issue, we invented a behavioral procedure that allowed us to assess how macaques—the closest model organism to humans where neuronal population activities are monitored—assign value to probabilistic predictions of novelty and physical rewards, and to study how novelty and reward interact in value-guided choice. We then uncovered a temporal-orbital neural pathway that causally regulates novelty-reward interactions during value-based decision-making. The pathway comprises the anterior medial ventral cortex (AVMTC), the final stage in the ventral visual stream linked to the detection and expectation of novelty (Bussey et al., 2005; Bussey & Saksida, 2002; Li et al., 1993; Ogasawara et al., 2022; Xiang & Brown, 1998; Zhang et al., 2022; Zhu et al., 1995; Zhu & Brown, 1995), and the orbitofrontal cortex (OFC; area 13m), an area causally linked to reward value-based decision-making (Padoa-Schioppa & Cai, 2011; Padoa-Schioppa & Conen, 2017; Rolls, 2000; Wallis, 2007). As monkeys chose among familiar offers associated with varying expectations of novel objects and expectations of rewards, expectations of novel objects increased the preference for large juice reward. Neural activity in the AVMTC-OFC circuit reflected this novelty-reward interaction. Specifically, novelty expectation signals emerged earlier in AVMTC, whereas subjective value, a neural signal reflecting the expectation of both novelty and reward, was more prominently and stably encoded in OFC. Chemogenetic disruption of the AVMTC**→**OFC pathway altered the impact of expected novelty on valuation of physical reward. Hence, our results suggest that the ventral visual system regulates novelty-reward interactions during decisions through direct projections to the prefrontal cortex.

## RESULTS

### Novelty and reward interact guiding motivation and preference

Monkeys participated in a behavioral procedure that contained distinct contexts or *blocks of trials* in which expected novelty was associated with high or low reward value (Figure 1A, B). In block 1, a greater chance of future novelty was associated with a greater chance of receiving big juice rewards (“novel→good”). In block 2, a greater chance of novelty was associated with a greater chance of receiving small rewards (“novel→bad”). Specifically, in block 1, five familiar visual stimuli served as *offers* that predicted future novel versus familiar objects with five distinct probabilities (% chance of novel/familiar objects: 0/100, 25/75, 50/50, 75/25, 100/0). A novel object was always followed by a big reward while a familiar object was always followed by a small reward. Oppositely, in block 2, five distinct offers also predicted novel objects with the same five probabilities, but here, a novel object was always followed by a small reward while a familiar object was always followed by a big reward. Hence, offers associated with a greater probability of obtaining novel objects had greater reward value in block 1, while offers associated with a greater probability of novel objects had relatively low reward value in block 2. Additional control blocks (blocks 3-4; Figure 1A) contained offers that predicted rewards with different probabilities but contained no novelty (block 3; “no novelty”) and offers that predicted novelty with different probabilities but kept expected reward constant and equal to the other blocks (block 4; “novel→neutral”).

**Figure 1.**
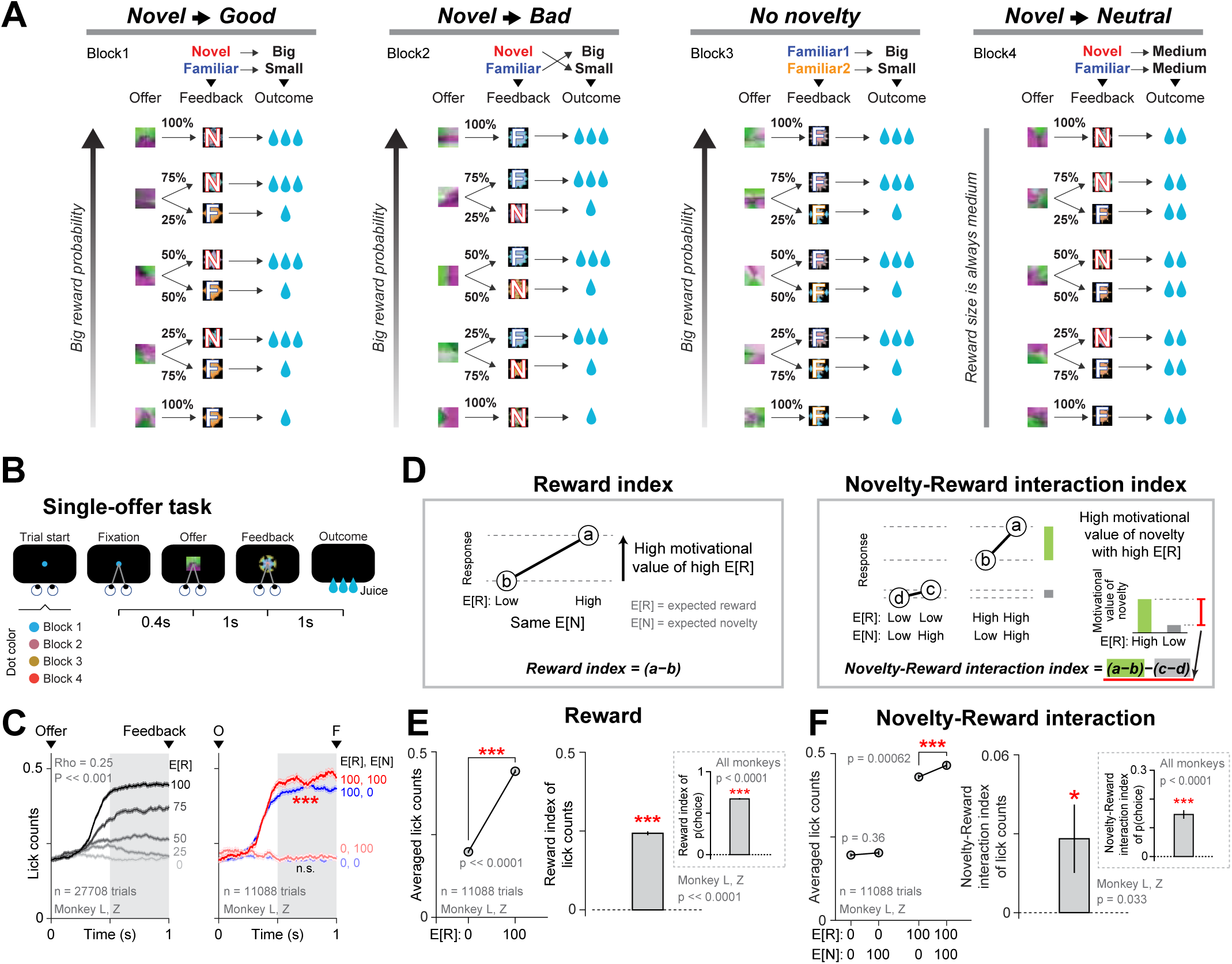
Behavioral interaction between physical reward and perceptual novelty. (**A**) Task conditions. **(B**) Single-offer task diagram. Each trial began with a central fixation dot. After gaze was maintained at the dot for 0.4s, the dot was replaced by an offer. The offer remained for 1s and was then replaced by a feedback object (novel or familiar). These objects were associated with large or small rewards as specified in **A** (also see Methods). To control for effects of reward and novelty on gaze, animals were required to keep fixation until juice reward. During blocked experiments in which the offers appeared in a block-wise fashion (as shown in **A**), the color of the initial fixation spot indicated the identity block (see Methods for further details). (**C**) Analyses of motivation with lick counts across all recording sessions (monkey L, n = 46 sessions; monkey Z, n = 37 sessions). **Left**: Shortly after offer onset, monkeys’ licking behavior scaled with expected reward associated with the probabilistic offers. Darker colors indicate higher expected reward (E[R]; probability of big reward). Spearman’s rank correlation result is indicated within the plot. The five different reward probabilities are indicated to the right of the time courses. **Right**: The licking reveals a motivational interaction between expected reward and novelty. Data is shown for high expected reward (E[R]) & high expected novelty (E[N]; probability of novelty) offers (red), high expected reward & low expected novelty offers (blue), low expected reward & high expected novelty offers (light red), low expected reward & low expected novelty offers (light blue). Animals showed a greater degree of offer presentation-related licking in high expected reward trials when expected novelty was also high (red vs. blue; P = 0.00062). In contrast, licking didn’t differ among low expected reward trials when expected novelty was low versus high (light red vs. light blue; P = 0.36). Error bars denote SEM. Asterisks indicate a significant difference (*** P < 0.001, Wilcoxon ranksum test). n.s. – non-significant. Gray areas indicate test windows. s – seconds. (**D**) Schematics of (**left**) Reward index quantifying the behavioral effect of expected reward (E[R]) and (**right**) Novelty-Reward interaction index quantifying the interaction of expected reward (E[R]) and expected novelty (E[N]). Reward index measures the difference in response to offers with high E[R] versus low E[R], keeping E[N] constant. Novelty-Reward interaction index measures the difference in response to E[N] when E[R] is high versus when E[R] is low. Further details about the indices are in Methods. (**E**) **Left**: Licking in response to high vs. low E[R] offers (big reward versus small reward). Error bars represent SEM. Asterisks indicate a significant difference (*** P < 0.001, Wilcoxon ranksum test). **Right**: Reward index calculated for the same data. Asterisks denote a significant difference (*** P < 0.001, Wilcoxon ranksum test). **Inset**: Reward index computed from monkeys’ choices instead of licks (Methods). The results are similar. P values are indicated on plots. Asterisks denote a significant difference (*** P < 0.001, bootstrapping with 10,000 repetitions). Error bars represent SD. (**F**) **Left**: Licking response (same trial types as in **C**-right). Conventions are the same. **Right**: Novelty-Reward interaction index of lick counts. * P < 0.05, permutation test with 10,000 repetitions. **Inset**: Novelty-Reward interaction index of choice preference. Consistent with the same index of lick counts, the Novelty-Reward interaction index of choice preference was significantly higher than zero. P values are reported on plots. *** P < 0.001, bootstrapping with 10,000 repetitions. Error bars represent SD.

The monkeys readily learned the meaning of the offers. This was verified with two behavioral measures. Following the presentation of the offers in the single-offer trials (Figure 1B), their anticipatory licking (Monosov et al., 2015) scaled with the probability of reward across (Figure 1C-left) and within all blocks that contained varying reward probability (blocks 1-3; Supplementary Figure 1). Also, when presented with choice trials (Methods), monkeys’ choice preferences co-varied with reward probability, indicating that they understood the meanings of the offers (Supplementary Figure 2-3).

We next assessed whether and how the expectation of novelty impacts the evaluation of expected juice reward. First, we concentrated on anticipatory licking – a well-accepted measure of motivational value. We asked: do monkeys lick more in anticipation of an increasing probability of physical juice reward when those rewards are also associated with the delivery of future visual novel objects? Figure 1C-right shows licking behavior for four key offers that allow us to qualitatively visualize the answer to this question and quantitatively assess the strength of the interaction between novelty and reward during offer evaluation: (trial type *a*; Figure 1D-right) a high chance of receiving a big reward and a novel object, (trial type *b*) a high chance of receiving a big reward and a low chance of receiving a novel object, (trial type *c*) a low chance of receiving a big reward and a high chance of receiving a novel object, and (trial type *d*) a low chance of receiving a big reward and a low chance of receiving a novel object. If novelty impacts reward expectation through an *additive* mechanism, we expect licking to be greater in trial type *a* than in trial type *b*, and greater in trial type *c* than in trial type *d*. If the mechanism is not purely additive, particularly displaying features of *multiplicative or gain-like* modulation, we would expect licking to be greater in trial type *a* than in trial type *b*, and roughly similar in trial type *c* than in trial type *d*. Indeed, licking behavior was significantly boosted when monkeys expected a big reward with a novel object (compare trial type *a* and *b*), but largely unaffected when monkeys expected a small reward with a novel object (compare trial type *c* and *d*) (Figure 1C, right; quantifications are shown in Figure 1F-left and Supplementary Figure 4). This suggests that when comparing motivated behavior across blocks 1 and 2, the effect of novelty on reward probability-driven motivation was relatively more multiplicative.

To further assess this effect, we quantified the magnitude of the impact of expected reward and the novelty-reward interaction on behavior with indices based on trial-by-trial licking (schematics in Figures 1D; details in Methods). Consistent with the data in Figure 1C, we found that as monkeys displayed anticipatory licking, they displayed both a positive Reward index and a positive Novelty-Reward interaction index (Figure 1E-F, right). Importantly, the same indices constructed from monkeys’ choice preferences also revealed positive and significant Reward and Novelty-Reward interaction indices (All monkeys together in Figure 1E-F, inset; each monkey separately in Supplementary Figure 5A). Thus, monkeys consistently chose offers more with high expected reward and high expected novelty over offers with high expected reward and low expected novelty.

We verified these results with a model-based approach across all trial types, enabling us to evaluate the magnitude and direction of key attribute effects underlying complex choice patterns across twenty distinct offers (as in Supplementary Figure 3). A generalized linear model (GLM) assessed the relationship between monkeys’ choice preference and offer attributes, namely expected reward (E[R]; probability of big reward), expected novelty (E[N]; probability of novelty), and their interaction (E[N] x E[R]). This analysis further confirmed that the effect of expected reward (E[R]) and the interaction between novelty and reward (E[N] x E[R]) were both positive and significant in the choice behavior across all monkeys and all trial types (Supplementary Figures 6A). This result remained robust even after restricting the analysis to offers from the key blocks that manipulated the novelty-reward contingency (blocks 1 and 2; statistical values are described in Supplementary Figure 6A legend), and even after adding a control regressor to the model indicating the possibility of gaining reward from those key offers (Methods; Supplementary Figure 6B).

We also observed an intriguing tendency for animals to behaviorally prefer offers from blocks where novelty and reward interacted. That is, animals generally chose to gain reward from offers that originated from blocks 1 and 2 (where reward was contingent on the presence or absence of novelty) in preference to the corresponding offers from block 3 (where reward was not related to the presence or absence of novelty; Supplementary Figure 2-inset). Importantly, this preference was distinct from the novelty-reward interaction or from a simple preference for novelty, and they could be measured with separate terms in the GLM (Methods; Supplementary Figure 6B).

In summary, the monkeys’ behavior indicated that they displayed a greater preference for objects that predicted *a greater probability of larger physical reward when they also predicted a greater probability of future perceptual novelty*. In other words, expectations of future novelty boosted the motivational value of reward-predicting objects.

We next investigated the neural circuits that might give rise to this modulation of reward-guided behavior by novelty. An important candidate is the anterior ventral medial temporal cortex (AVMTC), which includes the perirhinal cortex and medial IT (area TEav). Many AVMTC neurons strongly discriminate between novel and familiar objects (Li et al., 1993; Ogasawara et al., 2022; Tamura et al., 2017; Xiang & Brown, 1998; Zhang et al., 2022; Zhu et al., 1995; Zhu & Brown, 1995) and selectively respond to familiar visual stimuli that predict future novel objects (Ogasawara et al., 2022). The AVMTC has strong reciprocal connections with the orbitofrontal cortex (OFC; area 13m; Van Hoesen, Pandya, and Butters 1975; Morecraft, Geula, and Mesulam 1992; Barbas 1993; Suzuki and Amaral 1994; Carmichael and Price 1995), a brain area linked to many reward-guided decision-making-related processes (Padoa-Schioppa & Cai, 2011; Padoa-Schioppa & Conen, 2017; Rolls, 2000; Rushworth et al., 2011; Wallis, 2007), and their interactions have been implicated in valuation and decision-making computations that require visual information and associative memory (Clark et al., 2013; M. A. Eldridge et al., 2021; M. A. G. Eldridge et al., 2016; Monosov, 2024; Pelletier et al., 2021; Pelletier & Fellows, 2021). To assess how AVMTC-OFC circuit contributes to novelty-reward decisions, we first recorded neurons in these regions while monkeys participated in our behavioral task (Figure 2) and then applied the insights from those experiments to generate and test hypotheses through pathway-specific chemogenetic circuit manipulation (Figures 3-4).

**Figure 2.**
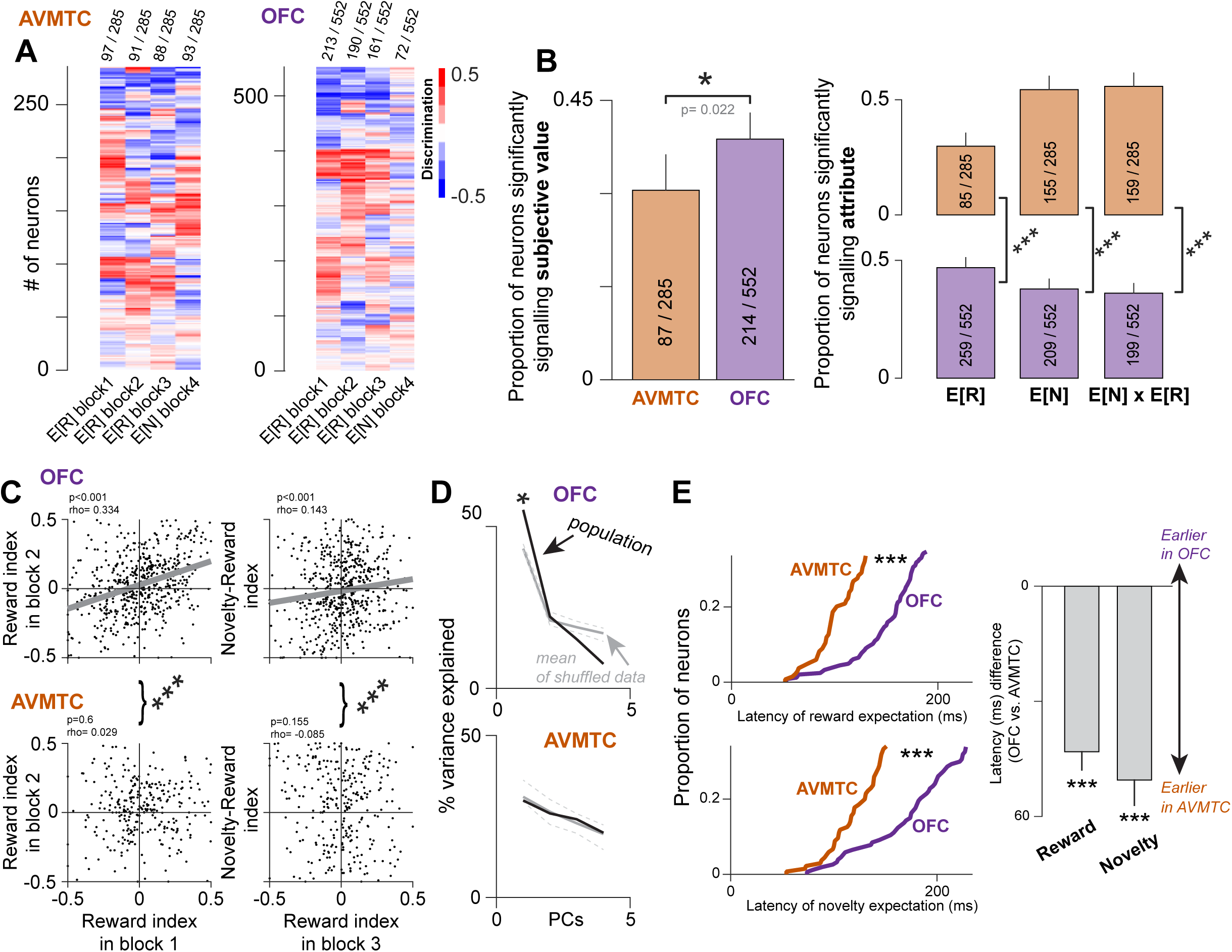
AVMTC and OFC reflect physical reward, object novelty, and novelty-reward interaction. (**A**) Heatmaps of single neurons (rows) selectivity quantified by discrimination of expected reward in blocks 1-3 (first three columns in each heatmap) and expected novelty in block 4 (last column in each heatmap). Values greater than 0 (red) indicate higher activity for a greater E[R] in the first three columns, and a greater E[N] in the last column. Values less than 0 (blue) indicate higher activity for a lower E[R] in the first three columns, and a lower E[N] in the last column (see Methods for details). Numbers of significant neurons in each column are indicated above. (**B-left**) Proportion of neurons in AVMTC and OFC that significantly scaled their activity with subjective value (Spearman’s correlation; P < 0.05). Subjective value was derived from monkeys’ choice behavior using the same GLM models also used for behavioral analyses in Figure 1 (Methods). OFC was more enriched with subjective value-related neurons (P < 0.001). (**B-right**) Proportion of neurons encoding expected reward (E[R]), expected novelty (E[N]), and their interaction (E[N] x E[R]) in each area. Each attribute was significantly different (P < 0.001) across areas: OFC was significantly enriched in E[R] relative to AVMTC, while AVMTC was more enriched in E[R] and E[N] x E[R]. (**C-left**) Correlations of ROC-based indices measuring expected reward coding (Reward indices) in blocks 1 and 2 were significant in OFC but not AVMTC (Spearman’s correlations; P < 0.001 in OFC, P > 0.05 in AVMTC). These correlations were significantly different (indicated as asterisks between upper and lower plots; ***P < 0.001; permutation test). (**C-right**) An ROC-based index measuring the modulation of big-reward expectation signals by novelty obtained from blocks 1 and 2 (Novelty-Reward index) was correlated with an independent Reward index (computed from trials in block 3) in OFC but not AVMTC. Conventions and tests are the same as on the left. (**D**) Dimensionality reduction by principal component (PC) analysis of single neuron data in A. PCs on x-axis, and % variance explained on y-axis. Gray lines show the maximum, mean, and minimum variance explained by each component in shuffled data (n=1000 shuffles; during each shuffle the elements in each row, in other words within each neuron, are shuffled). An asterisk indicates variance explained above the max of the shuffled data. (**E**) Reward and novelty expectation signals emerge earlier in AVMTC than OFC (P < 0.001). Cumulative distributions for single neuron latencies shown on the left and latency differences on the right. ms – milliseconds.

**Figure 3.**
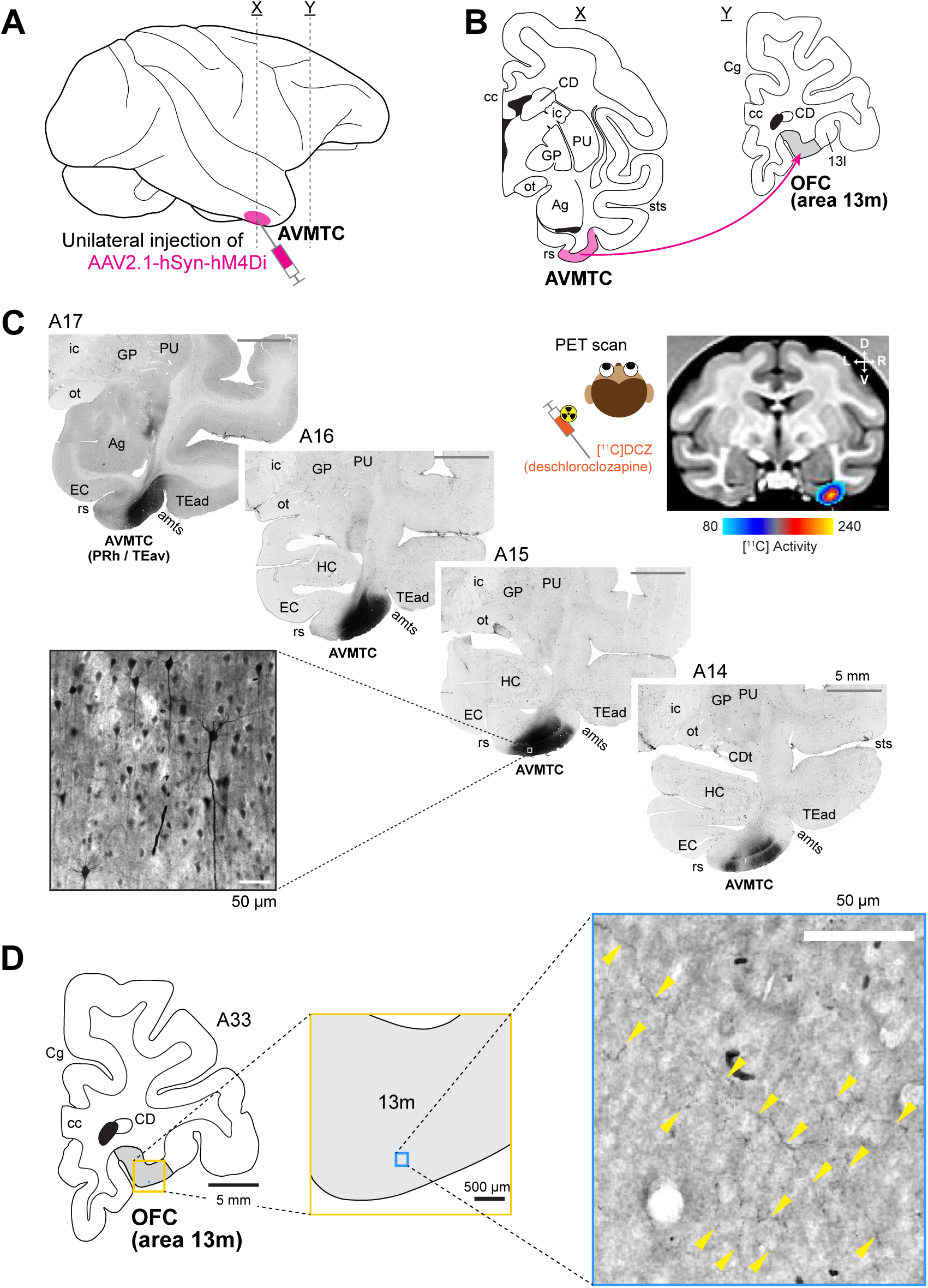
*Validation of viral expression for the disruption of the* AVMTC → OFC *pathway*. (**A-B**) Experimental plan. Following unilateral injection of AAV2.1-hSyn-hM4Di-GFP into AVMTC (**A**), we aim to disrupt the AVMTC → OFC pathway by activating the inhibitory DREADD (hM4Di) on AVMTC axons within OFC (**B**). Representative coronal planes correspond to the vertical lines in the lateral view (<u>X</u> and <u>Y</u> in 3A). Ag amygdala; CD caudate; Cg cingulate cortex; GP globus pallidus; PU putamen; cc corpus callosum; ic internal capsule; ot optic track; rs rhinal sulcus; sts superior temporal sulcus. (**C**) Immunohistochemistry visualizing green fluorescent protein (GFP) expression at the virus injection sites within AVMTC shown on four representative brain slices from Monkey Z. The approximate rostrocaudal distance (millimeters) from the interaural line is indicated at the top of each image. CDt caudate tail; HC hippocampus; PRh perirhinal cortex; TEad dorsal subregion of anterior TE; TEav ventral subregion of anterior TE; amts anterior middle temporal sulcus. Other abbreviations are described in 3B. **Left inset:** A magnified image of labeled AVMTC neurons at the injection site. **Right inset:** PET scan utilizing radioactive deschloroclozapine ([^11^C] DCZ) – a molecule that selectively binds hM4Di - visualized in vivo expression of hM4Di in AVMTC in the same monkey. MRI coronal slice at the level of AVMTC shows specific PET-assessed uptake of [^11^C] DCZ at the injection site. Color represents PET activity averaged over 90 -120 minutes following injection of [^11^C] DCZ (also see Supplementary Figure 11 and Methods for further details). D dorsal; V ventral; L left; R right. (**D**) Immunohistochemistry revealed GFP-labeled AVMTC axon-terminals in area 13m. **Left:** Illustration of a representative coronal section where dense axon-terminals were observed in area 13m. The gray area represents area 13m. The approximate rostrocaudal distance from the interaural line is at the upper right. **Middle:** A magnified illustration of area 13m corresponding to the yellow square in 3D-left. **Right:** A highly magnified image of area 13m corresponding to the blue square in 3D-middle. Most of the labeled axon terminals are indicated by yellow arrows.

**Figure 4.**
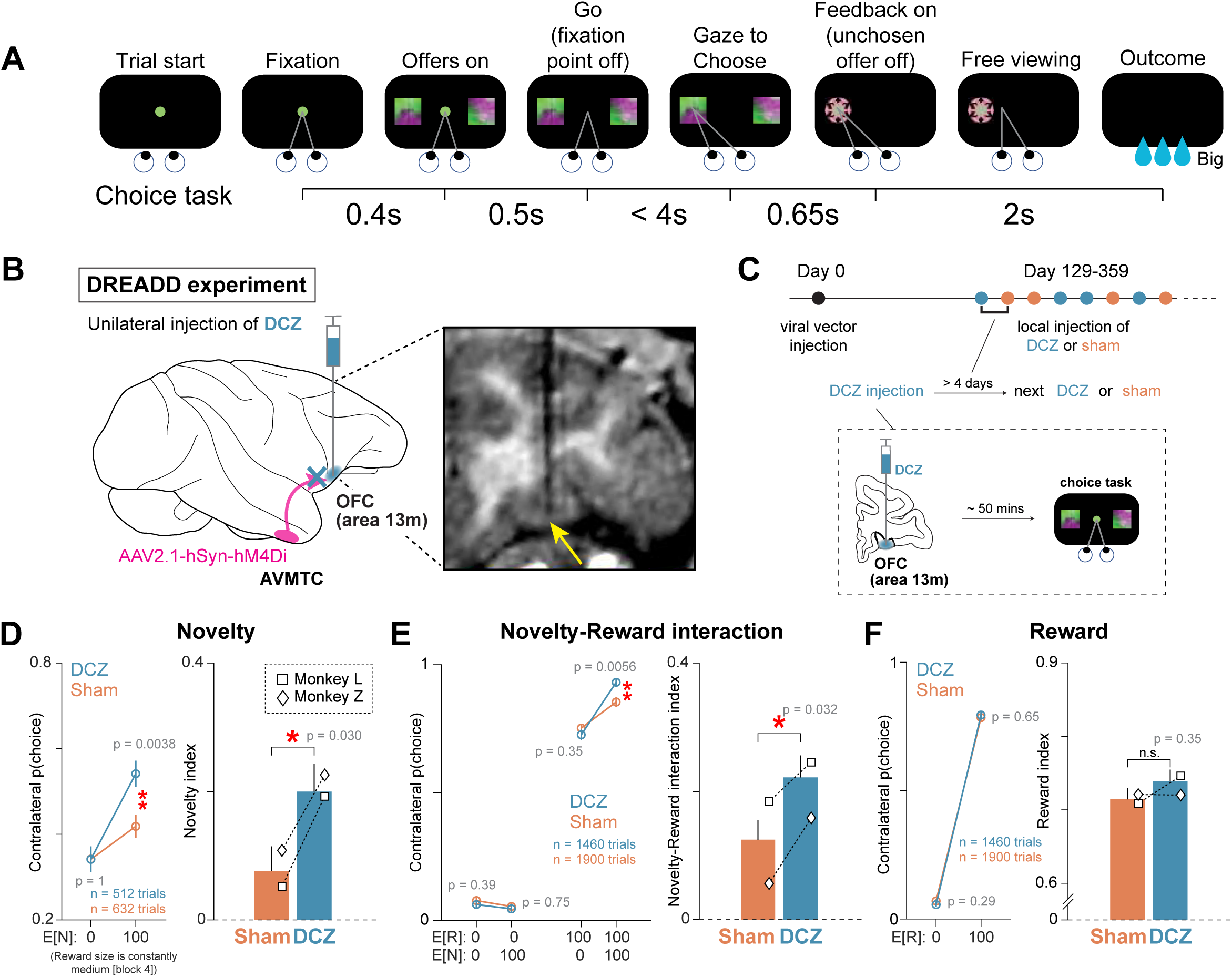
*Chemogenetic circuit manipulation reveals the* AVMTC → OFC *pathway causally impacts the novelty-reward interaction.* (**A**) Schematic diagram of a choice task. Two offers were randomly selected from twenty offers (Figure 1), and the animal chose among them. (**B**) **Left:** Illustration of chemogenetic unilateral disruption of AVMTC→OFC pathway by local DCZ infusion into area 13m. **Right:** The drug was injected into the sites where value-related neurons were enriched in area 13m. A representative MR image with a tungsten electrode (the tip is indicated by a yellow arrow) in one such site in area 13m. (**C**) Longitudinal schematic diagram of the long timeline of our DREADD experiments. DCZ (blue) and sham (orange) sessions were mixed to prevent animals’ ability to predict the nature of the experiment with a minimum of four days from a DCZ session to the next DCZ or sham session (Methods). (**D-F**) Comparing DCZ (blue) and sham (orange) sessions indicated that DCZ-mediated AVMTC → OFC disruption altered the impact of novelty on decision-making. (**D**) Choice rates for contralateral offers with high vs. low E[N] in block 4 where reward size was constantly medium (left) and Novelty index obtained from the data on the left (right; see Methods for its calculation). (**E**) Choice rates for the four contralateral offer types as in Figure 1C-right (left) and the associated Novelty-Reward interaction index (right). (**F**) Choice rates for contralateral offers with high vs. low E[R] (left) and the associated Reward index (right). Data across all sessions from the two injected monkeys was used for analyses (DCZ: 7 and 6 sessions; sham: 5 and 6 sessions for monkeys L and Z, respectively). The same calculations for ipsilateral offer presentations in Supplementary Figure 13 show no significant ipsilateral effects. Bars denote indices across all monkeys; results for each monkey separately are shown by different symbols (square, monkey L; diamond, monkey Z). Contralateral Novelty and Novelty-Reward interaction indices increased following DCZ injection relative to sham, indicating an alteration of the impact of novelty on the decision process (see GLM-based analyses showing similar results in Supplementary Figure 14). Notably, Reward index did not change, suggesting that disruption of AVMTC inputs does not alter the valuation of expected reward. Error bars for choice percentages and indices indicate SEM and SD, respectively. Asterisks indicate a significant difference (* P < 0.05, ** P < 0.01, *** P < 0.001, permutation test with 10,000 repetitions). n.s. – non-significant.

### Comparing reward and novelty signals in AVMTC and OFC

Both OFC and AVMTC contained a significant number of neurons whose responses correlated with reward and novelty expectation (Figure 2A). We, therefore, applied theory-guided analyses to assess their similarities and differences. These analyses together suggested that OFC was more closely tied to representing the subjective value or preference for reward and novelty-predicting offers than AVMTC, while AVMTC was relatively more dedicated to processing novelty and novelty expectation.

For the first analysis, we used a standard model-based approach to derive a quantitative measure of the subjective value of each offer based on the monkeys’ choices (Methods). We then correlated this value with each neuron’s responses to the offers’ presentations (Bromberg-

Martin et al., 2024). Relative to AVMTC, more OFC neurons’ responses were significantly correlated with subjective value (Figure 2B-left; correlation threshold: P < 0.05; P = 0.022). Also, like the monkeys’ choices (Supplementary Figure 6), OFC neurons more commonly reflected the probability of physical reward, while AVMTC more frequently reflected the probability of future novelty and the novelty-reward interaction (Figure 2B-right). Both areas contained significant numbers of neurons that signaled E[N] x E[R] though AVMTC was significantly more enriched (Figure 2B-right; also see Supplementary Figure 7).

If OFC is more closely tied to guiding behavior based on the subjective value of visual objects, it ought to signal expected physical reward relatively consistently across contexts or blocks (following the monkeys’ stable preference for higher physical reward across the task blocks; Supplementary Figure 6). Indeed, qualitative inspections of the heatmaps in Figure 2A suggest that expected reward is encoded relatively more stably in OFC than in AVMTC. In OFC, neurons signaling value positively (that is, with more spikes in response to the prediction of big reward; *red*) often consistently did so across blocks, and neurons signaling value negatively (*blue*) tend to also do this consistently across blocks 1-3 (first three columns in heatmaps in Figure 2A). In contrast, AVMTC neurons tended to scale with reward expectation in a relatively more “disorganized” manner, for example, in one block but not in others. Indeed, as suggested by these observations, further analyses revealed that during offer presentation, reward expectation signals were correlated across blocks more strongly in OFC than AVMTC (Figure 2C-left; OFC, rho = 0.334, P < 0.001; AVMTC, rho = 0.029, P = 0.6; OFC greater than AVMTC, P < 0.001, permutation test). The results held when other blocks were considered (Supplemental Figure 8). Thus, reward probability, a key decision variable mediating monkeys’ preference consistently across the task blocks (Supplementary Figure 6), was relatively more consistently represented by OFC neurons across the task blocks or contexts.

If OFC is indeed more closely tied to subjective value than AVMTC, then OFC ought to more closely reflect monkeys motivation by both expected reward and by the novelty-reward interaction (Figure 1C-F, Supplementary Figures 4, 5A, and 6). This is what we observed (Figure 2C-right). Across the population, OFC neurons had a stronger positive correlation between the index that measured expected reward coding and the index that measured the modulation of big-reward expectation by novelty (OFC, rho = 0.143, P < 0.001; AVMTC, rho = - 0.085, P = 0.155; the correlations were significantly different across areas, P < 0.001; permutation test; Methods). Thus, OFC neurons sensitive to expected physical reward tended to be sensitive in the same manner to the interaction of physical reward and novelty.

By contrast to OFC and to monkeys’ preferences for expected reward across the block contexts, AVMTC did not have a consistent representation of physical reward value across block contexts. Principal component (PC) analyses of neural response patterns in the heatmap in Figure 2A revealed that much of OFC response patterns were explained by a single PC, while AVMTC response patterns had lower percentage variance explained by PCs in a manner that was similar for actual vs. shuffled data (n=1000 shuffles; Figure 2D; also see Supplementary Figure 9).

What then is the role of AVMTC? We reasoned that AVMTC may be a crucial source of input, where distinct reward and novelty signals emerge at shorter latencies and are then sent to be integrated in the OFC. However, another possibility could be that these signals emerge earlier in OFC and are then sent to AVMTC to modulate ongoing sensory and memory processing. Our data provide evidence for the first scenario. The latencies of reward and novelty expectation signals were significantly earlier in AVMTC than in OFC (P < 0.001; Figure 2E; also see Supplementary Figure 10). Like these main effects, a GLM based analysis (same as in Supplementary Figure 10) revealed that the interactions between reward and novelty were also detectable in AVMTC before OFC (P < 0.01 for either GLM with or without block effect).

In summary, OFC contained a relatively consistent (across-block) representation of physical reward value more clearly aligned with the monkeys’ preferences, suggesting that it may be downstream of AVMTC in value-based decision computations. AVMTC was relatively more dedicated to processing novelty and is well positioned to provide necessary input to OFC to support value-related computations and to guide novelty- and reward-guided behavior.

### Chemogenetic disruption of the AVMTC→OFC circuit during novelty-reward interactions

Our data thus far suggest that AVMTC could broadcast novelty expectation signals, including to OFC, and that OFC is more closely tied to integrating such novelty signals and reward to guide behaviors. Our results from the following experiment – a projection-specific chemogenetic disruption of the AVMTC-OFC circuit – strongly support this hypothesis (Figure 3, 4).

The logic of the circuit disruption is as follows. We sought to dampen the effect of AVMTC inputs on OFC computation as monkeys made decisions about future reward and novelty. To do so, we first unilaterally expressed inhibitory designer receptors exclusively activated by designer drugs (DREADDs; hM4Di) in the right AVMTC through the injection of an adeno-associated viral vector (AAV2.1-hSyn-hM4Di-GFP; Figure 3A, Supplementary Figure 11F; Methods). AAV2.1-hSyn produces robust protein expression on axon terminals and has been used to modulate the impact of axon terminals on behavior in non-human primates (Kimura et al., 2023; Nagai et al., 2024; Oyama et al., 2021). To manipulate the circuit, we then injected the DREADD agonist (Deschloroclozapine [DCZ]; Nagai et al. 2020) unilaterally into the right OFC, the site of the AVMTC projections (Figure 3B). We then quantified whether and how reward- and novelty-guided decision-making changed.

We verified the efficacy of viral transfection in AVMTC both in vivo and ex vivo. First, we tagged DCZ, a high-affinity and highly selective agonist binding hM4Di, with radioactive ^11^C (Tian et al., 2015) and conducted PET imaging with [^11^C]DCZ to visualize in vivo hM4Di expression in the two monkeys (Figure 3C-right inset; see Supplementary Figure 11A-B top for each monkey). [^11^C]DCZ showed potent and selective binding in the injected region within AVMTC, demonstrating successful transfection of hM4Di (raw data and time course are shown in Supplementary Figure 11A-D). Second, we further confirmed the expression at the level of cell bodies and their axon boutons using immunohistochemistry. The AVMTC contained large numbers of transfected neurons, as shown in Figure 3C-left inset. Notably, labeled axon boutons were also found within OFC, area 13m (Figure 3D-right, yellow arrows) of the same hemisphere as the injected AVMTC. This confirmed that AAV2.1-hSyn can be used for transfections at the site of axon terminals in primate cortico-cortical circuits. This area 13m is where we (as shown in Figure 2) and others found many value-related neurons (Duuren et al., 2008, 2009; Feierstein et al., 2006; Padoa-Schioppa & Assad, 2006; Roesch & Olson, 2004; Roitman & Roitman, 2010; Rolls & Baylis, 1994; Schoenbaum et al., 1998; Schoenbaum & Eichenbaum, 1995; Sul et al., 2010; Thorpe et al., 1983; Tremblay & Schultz, 1999; Wallis & Miller, 2003) and where the DCZ injections were made in subsequent experiments (Figure 4B).

To measure the impact of DCZ-mediated disruption of AVMTC-OFC signaling on value-guided decision-making, we let the monkeys choose among pairs of offers drawn from the full set of twenty possible offers in the task (Figures 1A, 4A, Methods). We then compared behavioral performance on DCZ versus sham sessions (Figure 4B-C; also see Supplementary Figure 12) using behavioral indices for reward, novelty, and the novelty-reward interaction (Figure 1D, Methods) separately for offers presented contralaterally or ipsilaterally to the injection site (Methods).

We found that the circuit disruption altered the impact of novelty on choice in a spatially dependent manner (DCZ, n = 4929 trials; sham, n = 6365 trials; Figure 4D-F). Both Novelty and Novelty-Reward interaction indices significantly increased on the contralateral side (Figure 4D-E, right). The increase in the Novelty index was caused by an increase in the preference for offers associated with higher expected novelty in block 4, where novelty had no extrinsic reward value (P = 0.0038, Figure 4D-left). The increase in the Novelty-Reward interaction index was caused by an increase in the choice of the offer with high expected reward and novelty, increasing the gain-like interaction between expected reward and novelty (P = 0.0056, Figure 4E-left). This effect was spatially dependent: the indices for ipsilateral choice preference were not changed by DCZ (Supplementary Figure 13-left and middle). This effect was also specific to novelty and its interaction with reward, not to reward *per se*. The Reward index did not change for either contralateral or ipsilateral offers. This suggests that DCZ effects were not simply due to changes in object-value memory, perception, or general reward-guided motivation (Figure 4F, Supplementary Figure 13-right).

To validate these results, we next turned to GLM-based analyses that considered all choice trials across all offers, incorporating new regressors for the visual hemifield of chosen offers (contralateral or ipsilateral) and drug (DCZ or sham). The results, like the behavioral indices, show that the novelty-reward interaction increased for contralateral relative to ipsilateral offers (increased effect of contra x E[N] x E[R] in trial-based and session-based analysis [DCZ, n = 13; sham, n = 11 sessions], Supplementary Figure 14, 15).

Finally, to verify whether DCZ injection had any off-target effects on behavior, we conducted unilateral local injections of DCZ into the same brain area, area 13m, but in the opposite hemisphere from virus-injected AVMTC. There were no significant changes in overall choice probabilities, indices, or GLM results (Supplementary Figure 16).

In sum, these results show that the disruption of the AVMTC→OFC pathway could alter the impact of novelty expectation on the valuation of expected rewards during value-based decision-making.

## DISCUSSION

Context-dependent valuation and integration of novelty and physical reward value are key aspects of human and animal cognition and behavior. We provide strong evidence that an AVMTC-OFC circuit causally regulates this process, with each area contributing in distinct manners.

AVMTC is relatively more enriched with novelty-expecting and familiarity-expecting neurons, while OFC is more enriched with reward-expecting neurons (Figure 2B-right). Reward-related neurons in OFC often reflected the interaction of physical reward and novelty that guided subjective value and motivated behavior (Figure 2C), and, as a population, OFC neurons resembled monkeys’ behavioral preferences (Figure 2B). These and other results from our study suggest that OFC could regulate value-based choice across reward and novelty in cooperation with AVMTC. We provided causal evidence for this hypothesis by directly disrupting the functional state of the AVMTC projection to OFC. We showed that this disruption altered the impact of novelty on reward probability-driven behavior. Therefore, AVMTC – a brain region previously associated with novelty detection and memory (Brown & Aggleton, 2001; Miyashita, 2019; Monosov, 2024; Murray & Richmond, 2001; Suzuki & Naya, 2014) – also participates in modulating the valuation of physical reward in response to novelty expectations through projections to the OFC in manners that include gain-like interactions (Figure 4E).

Current computational theories, as well as artificial intelligence benchmarking, suggest that curiosity-related learning and context-dependent behavioral control may depend on separate processing of physical extrinsic reward value and an intrinsic reward of novelty (Gottlieb et al., 2016; Y. Kim et al., 2019; Modirshanechi, Kondrakiewicz, et al., 2023; Modirshanechi, Lin, et al., 2023; H. A. Xu et al., 2021). This architecture could result in separate action planning strategies or policies, which then could be combined downstream, for example, to control learning and valuation through the integration of extrinsic and intrinsic reward (Aubret et al., 2023; Bellemare et al., 2016; Burda et al., 2018; Gottlieb et al., 2016; Houillon et al., 2013; Jaegle et al., 2019; Kakade & Dayan, 2002; Modirshanechi, Kondrakiewicz, et al., 2023; Savinov et al., 2019; Tang et al., 2017). Indeed, some recent experimental evidence supports that distinct neural pathways exist for reward-driven and novelty-driven behaviors (Ahmadlou et al., 2021; Menegas et al., 2017; Ogasawara et al., 2022), and others support that the modulation of reward-related signals by novelty change during learning as novel stimuli become familiar (Tremblay et al., 1998; Lak et al., 2016). However, how novelty and reward expectations are integrated in economic choice to guide behavior or neural activity has remained poorly understood. Our current results indicate that single neurons in an area known to mediate physical reward-guided decisions (OFC) reflect novelty-reward interactions, suggesting that novelty can directly impact computations closely tied to generating decision preference and value (Aubret et al., 2023; Bellemare et al., 2016; Burda et al., 2018; Gottlieb et al., 2016; Houillon et al., 2013; Jaegle et al., 2019; Kakade & Dayan, 2002; Modirshanechi, Kondrakiewicz, et al., 2023; Savinov et al., 2019; Tang et al., 2017). Therefore, the brain may have the mechanisms to control behavior either through distinct or integrated reward and novelty computations, depending on task demand and context (Monosov, 2024).

Animals’ physical reward probability-driven behavior remained intact despite the pathway disruption. This suggests that the increased influence of novelty expectation caused by the AVMTC→OFC pathway disruption is not simply due to changes in perception or object-reward associations. This fits well with studies in primates that showed that perirhinal cortex ablation does not impair perception or the retention of reward associations during decisions (Easton & Gaffan, 2000; M. A. Eldridge et al., 2018; Thornton et al., 1998). Instead, the lesion of area TE (a region lower in the putative ventral visual pathway hierarchy; Bussey and Saksida 2002; Bussey et al. 2005) impairs perception (M. A. Eldridge et al., 2018) and dissection of this area with the ventral prefrontal cortex impairs the object-reward associations (Parker & Gaffan, 1998).

A previous paper found that while expected reward amount associated with visual stimuli can be decoded earlier in temporal cortex (area TE), the dorsolateral prefrontal cortex signaled the expected reward value of objects relatively more independently of high-level object features (Sasikumar et al., 2018). This fits well with our results (Figure 2) and highlights a circuit architecture in which object information is further abstracted at the prefrontal cortex (Freedman et al., 2001, 2003; McKee et al., 2014)

Disrupting AVMTC inputs to OFC increased the impact of novelty expectation on object value-related preference (Figure 4). Eldridge and his colleagues (2016) show that, in a task that measured object-reward-driven motivation (but not preference), disruption of perirhinal/rhinal→OFC pathway reduced error rates during low-value trials. However, Clark (2013) showed that permanent temporal–orbitofrontal disconnection lesions resulted in changes in object-value related motivated behavior that was due to factors beyond reward associative learning. While these prior instrumental paradigms differ from our novelty-reward choice task, and do not measure choice-preference or the impact of novelty expectation on behavior, their findings and our current data together point to a complex picture where AVMTC and OFC play multiple context-dependent roles in behavioral control.

AVMTC disruption may alter object recognition (e.g., overall increasing the novelty of visual objects), reduce the accuracy of the classification of stimuli into novel or familiar, and/or impact the effects of these processes on novelty or value related internal states (e.g., the subjective experience or attentional prioritization in response to novelty or familiarity). Also, in the current study, AVMTC contained subpopulations of neurons that either elevated or reduced their activity during novelty expectation. This contrasts with results from a previous task in which novelty was completely unrelated to reward outcomes and hence could not interact with or impact physical reward value or rate. There, AVMTC neurons tended to often be selectively excited by novelty expectations more than by familiarity expectations (Ogasawara et al., 2022). These observations imply that representations in AVMTC and their impact on OFC may be context-dependent, adapted to the novelty and reward statistics of the task at hand - a feature that could be a key factor for adaptive behavioral control in complex and volatile environments (for review see Monosov et al., 2022). In tasks in which reward and novelty interact, AVMTC may not always enhance the impact of novelty expectation on reward and could even reduce it.

OFC value computations influence animals’ preference and motivation (Ballesta et al., 2020; Elston & Wallis, 2025; Knudsen & Wallis, 2022; Wilson et al., 2014) and top-down attention (Hunt et al., 2018; Lim et al., 2011; McGinty et al., 2016; Perkins & Rich, 2024; Xie et al., 2018). When OFC computations are disrupted by reducing AVMTC inputs, other circuits, and brain areas involved in novelty detection and novelty seeking (Burns et al., 1996; Chen et al., 2020; Costa et al., 2014, 2019; Monosov et al., 2022; Ogasawara et al., 2022; Tapper & Molas, 2020; Yamamoto et al., 2012) may gain a relatively stronger influence over choice (particularly if AVMTC is not acting simply to increase the value of reward expectations when novelty is expected). For example, a faster process that guides animals towards future novelty through the zona incerta-brainstem circuits (Ahmadlou et al., 2021; Ogasawara et al., 2022) may become more dominant following the disruption of OFC.

These hypotheses are in line with previous proposals that novelty-seeking may be controlled by multiple circuits on multiple distinct time-scales allowing for faster- and slower-responses (relying on different levels of computational or processing complexity) to be adjusted to the context or environment at hand (Monosov et al., 2022). Future large-scale neurophysiological experiments are required to assess the validity of this proposal.

AVMTC inputs to OFC terminate preferentially in superficial layers in both primates and rodents (Agster & Burwell, 2009; Lavenex et al., 2002; Rempel-Clower & Barbas, 2000). Recent studies in rodents revealed mechanisms where inputs from higher areas to apical dendrites in cortical layer 1 control learning (Cichon & Gan, 2015; Doron et al., 2020; Lacefield et al., 2019; Leinweber et al., 2017), sensory perception (Manita et al., 2015; Takahashi et al., 2016, 2020; N. Xu et al., 2012), and adaptive behavior (Ranganathan et al., 2018), particularly by evoking Ca^2+^ signals in the dendrites of layer 5 pyramidal neurons. Therefore, studying the modulation of OFC dendritic activities by AVMTC inputs will provide a deep understanding of the mechanisms of how this pathway regulates object valuation.

How intrinsic motivation and physical reward interact to guide decision-making has been a mystery (Monosov, 2024). Our study shows that novelty can impact physical reward valuation and motivation in gain-like manners and that this effect may be sensitive to context or block (Supplemental Figures 2 and 12, insets). This regulation is supported by the AVMTC → OFC cortico-cortical circuitry, broadly demonstrating how the temporal visual system can directly impact motivation and valuation and providing a deeper understanding of and suggesting novel architectures for curious agents.

## SUPPLEMENTARY FIGURE LEGENDS

***Supplementary Figure 1. Raw offer-related licking behavior.*** Averaged lick counts from the test window, shown as a gray area in Figure 1C, are presented for each offer in each block (B) of trials (red, block1; blue, block2; green, block3; black, block4) for all and separately for each monkey. Correlation between lick counts and the probability of a big reward (E[R]) and novelty (E[N]) are shown for each block (Spearman’s rank correlations). Error bars denote SEM. Anticipatory licking scaled with E[R] within each block (blocks 1-3). In block 4, where reward probability was held constant (100% of medium reward), licking behavior did not scale with E[N] – consistent with our theory that novelty impacts licking through interaction with reward (see Figure 1). Licking data was collected across all recording sessions in all monkeys. Monkey A did not display offer-related licking during offer presentation; hence, it is not included in analyses of licking. This monkey readily performed choice trials and its data is therefore included in all analyses quantifying offer preference (measured by choice).

***Supplementary Figure 2. Raw choice behavior.*** Probability (P) of choosing each offer across the 4 blocks. Conventions are the same as in Supplementary Figure 1. Error bars denote SEM. Data was collected from all sessions during the period of neural recordings and sham sessions across all three monkeys (trial numbers for each block are displayed on the plots for all and separately for each monkey). In addition to the key result of our paper – the interaction of novelty and reward (Figure 1F), there were context-dependent effects of novelty. For example, offers in block 2, where novelty is associated with a small reward, tended to be less preferred than block 1 where novelty predicted a big reward (this was the case even for the 50% offers where expected novelty and expected reward were the same, but the associations were reversed [monkey L, P = 9.1 x 10^-12^; monkey Z, P = 1.0 x 10^-76^; monkey A, P = 0.0057, permutation test]). Also, offers in block 2 tended to be more preferred over offers in block 3, despite the matching expected reward. Due to these observations, we further confirmed our results throughout this paper by modeling the data with block effects in subsequent Supplementary Figures (also see Methods). **Inset**: Data of blocks 1-3 are overlaid.

***Supplementary Figure 3. Detailed choice preference matrices.*** (**A**) Pairwise choice probability (%) plots across all monkeys. The choice probability for each offer across blocks (B) 1-3. The choice probability for each offer in block 4 is shown on the right. Colormap denoting % choice is above. (**B**) Same data separately for each monkey.

***Supplementary Figure 4. The novelty-reward interaction in each monkey’s licking behavior.*** Data across all recording sessions for monkeys L and Z were included. Conventions are the same as in Figure 1F.

***Supplementary Figure 5. Reward, Novelty, and Novelty-Reward interaction indices.*** (**A**) Conventions are the same as in Figure 1E-F. **Insets**: showing choice preferences for conditions from which the novelty-reward interaction is calculated (same as in Figure 1F). Error bars denote SEM. * P < 0.05, *** P < 0.001, permutation test with 10,000 repetitions. (**B**) Novelty index of choice, quantifying the behavioral effect of the expectation of novelty (see Methods for its calculation). Amazingly, though Novelty index was negative in one monkey (monkey L), Novelty-Reward interaction index was positive across all monkeys, further highlighting one of the key points of our paper. Error bars denote SD. **P < 0.01, ***P < 0.001, bootstrapping with 10,000 repetitions. Trial numbers across animals are balanced when pooling for across animal index calculation.

***Supplementary Figure 6. GLM analyses of choice preference.*** (**A**) a GLM weights obtained from analyses of choice for E[R], E[N], and E[N] x E[R]. Data shown for all monkeys and for each monkey separately. All monkeys assigned significantly positive weights to the novelty-reward interaction (*E[N] x E[R]*). This was the case even after restricting the analysis to offers only from blocks 1 and 2 (monkey L, E[R]: P = 0, E[N] x E[R]: P = 5.4 x 10^-37^; monkey Z, E[R]: P = 7.7 x 10^-183^, E[N] x E[R]: P = 1.2 x 10^-9^; monkey A, E[R]: P = 2.0 x 10^-95^, E[N] x E[R]: P = 2.6 x 10^-5^, linear hypothesis test). (**B**) Another GLM model with a block-effect term produced similar results. Error bars indicate SEM. Asterisks represent a significant deviation from zero (***P < 0.001, linear hypothesis test).

***Supplementary Figure 7. Additional GLM analyses of neural data.*** (**A**) Same data as in Figure 2 modelled with a GLM with an added term for block effect (Methods). All formatting and conventions are the same. (**B**) Same GLM model as in Figure 2 fits data in the unblocked control version of the task (Methods). The key findings are similar. Novelty-related attributes are more prominently represented in AVMTC.

***Supplementary Figure 8. Correlation of reward value signals across blocks.*** Same format as Figure 2C. Reward indices obtained in block 1 (left) and block 2 (right) are correlated with it obtained in block 3. Within OFC, the correlations are significantly stronger (***P < 0.001; permutation test).

***Supplementary Figure 9. An additional assessment of clusterability of OFC and AVMTC.*** Data of each area (indicated by color) in Figure 2A was clustered by k-means with different cluster numbers (x-axis). Silhouette values, a measure clusterability (Methods), were consistently higher in OFC (filled circles indicate significance; P < 0.05, Wilcoxon rank sum test). The key point is that the values were higher in OFC regardless of the assumption about cluster number.

***Supplementary Figure 10. Latency analyses of reward and novelty expectation signals.*** (**A-B**) Data in the same format as Figure 2E-right. (**A**) Latency results are similar when we obtain brain area latencies using an ROC-based method or GLM-based methods with the standard model or one with an added block effect term (Methods). These results remain similar across blocked and unblocked versions of the task. (**B**) Latency results are similar across blocked and unblocked versions of the task (format is same as in Figure 2).

***Supplementary Figure 11. PET and immunohistochemistry.*** (**A-B**) PET data for each of the two monkeys. Sagittal (**left**) and coronal (**right**) slices at the level of AVMTC, showing specific uptake of [^11^C] DCZ at the viral injection site in monkey Z (**A**) and L (**B**). Specific uptake of [^11^C] DCZ largely confined to the injection site in AVMTC in both monkeys. **Upper:** Thresholded displays. **Lower:** Corresponding displays without thresholding. The MRI and PET data of both monkeys were registered to a T1-weighted macaque atlas representing 42 other animals. This atlas is shown as the underlay in **A-B**. P posterior, A anterior. Other conventions are the same as Figure 3C, right inset. (**C-D**) Binding over time. Temporal dynamics of the DCZ signal in monkeys Z (**C**) and L (**D**) over the interval 0 - 120 minutes following injection of [^11^C] DCZ. Red line, right AVMTC (injected side); blue line, left AVMTC (other side); black line, right putamen; grey line, right subcortical white matter. All time activity curves were extracted from 2.5mm radius spherical regions interest. Right AVMTC ROIs (red dot) were centered on the locus of peak activity. Left AVMTC ROIs (blue dot) were centered on the homotopic locus in the contralateral hemisphere. Putamen (black dot) and white matter (gray dot) ROI center coordinates were visually placed by reference to the co-registered atlas. After 20 minutes, the injected AVMTC had a greater DCZ signal than control regions, which persisted thereafter. Each plotted point represents decay-corrected PET activity summed over intervals of duration (4 - 10 minutes). (**E**) Photographs of the representative coronal section including area 13m (left), a magnification of area 13m (upper right), and a high magnification of axon terminals in area 13m (bottom right), corresponding to Figure 3D. All conventions are as in Figure 3D. (**F**) The sagittal (top) and coronal (bottom) MRIs with a tungsten electrode (indicated by yellow arrows) in AVMTC of monkey Z. PU putamen; HC hippocampus; CD caudate; Ag amygdala; ac anterior commissure.

***Supplementary Figure 12. Choice preference during DCZ pathway disruption experiments.*** (**A**) Choice preference for each offer across two monkeys, shown separately for contralateral (left) and ipsilateral offer presentation (right) relative to the drug-injected hemisphere. Red, block1; blue, block2; green, block3; black, block4. Darker and lighter colors in each panel represent data during DCZ injections versus sham, respectively. On the contralateral side, significant changes were observed in blocks 1 and 4. In block 1, preference for 100% reward / 100% novelty offers significantly increased following DCZ injection (P = 0.0056). In block 4, preference for the 100% novelty offer significantly increased (P = 0.0038). On the ipsilateral side, the significant changes were the reductions in choice preference for some of the novelty-predicting offers in Block 4 (75% and 50% expected novelty significantly decreased; P = 0.00034 and = 0.016, respectively). These results are quantified further by index-based analyses in Figure 4D-F for contralateral trials and Supplementary Figure 13 for ipsilateral trials. Error bars denote SEM. Asterisks indicate a significant difference in choice preference for DCZ versus sham trials (*P < 0.05, **P < 0.01, ***P < 0.001, permutation test). **Inset**: Data of blocks 1-3 are overlaid for DCZ injections (upper) and sham (bottom), respectively. (**B**) Plots and conventions are the same as in Supplementary Figure 3. **Left:** Pairwise choice probabilities in the sham sessions. **Middle:** Pairwise choice probabilities in DCZ injection sessions. **Right:** Differences in pairwise choice probabilities between DCZ and sham. The values represent the probability of each offer in the DCZ condition minus its corresponding probability in the sham condition, illustrating the change in choice preference by the pathway disruption. To statistically assess these data, we employed GLMs in Supplementary Figure 14-15.

***Supplementary Figure 13. Indices of ipsilateral choice preference in the pathway-disruption experiment.*** All conventions are as in Figure 4D-F.

***Supplementary Figure 14. GLM results for each monkey’s choice preference in the pathway-disruption experiments.*** (**A**) Similar to the results of index-based analysis in Figure 4D-E, the GLM showed that the unilateral DCZ injection into OFC changed the impact of novelty expectation on preference for expected reward for contralateral offers compared to ipsilateral ones (relative to the injection). To capture this spatial effect, we added the term of laterality (Contra) to the model in addition to the key attributes (E[R], E[N], and E[N] x E[R]; Methods). Fitted weights of the key attributes and the interaction between those attributes and the laterality (Contra) are shown for data across all DCZ (blue bars) and sham (orange bars) sessions. Significant changes were found in the interaction of the laterality and the novelty-reward interaction (contra x E[N] x E[R]) across monkeys. Error bars indicate SEM. Asterisks denote a significant deviation from zero (*P < 0.05, **P < 0.01, ***P < 0.001, Linear hypothesis test). (**B**) Effect of DCZ relative to sham in each session. To obtain these values, we subtracted the mean GLM weight from the sham sessions from single session weights in DCZ injections. White circles indicate medians. Each blue dot is each session’s results. Vertical black boxes indicate the interquartile range. Vertical black whiskers indicate a 1.5 x interquartile range. Violins represent kernel density estimates. Asterisks indicate a significant deviation from zero (*P < 0.05, Wilcoxon signed-rank test). Only key terms are shown in A-B (See methods for full terms used in the models).

***Supplementary Figure 15. GLM results with block-effect term in the pathway-disruption experiments.*** All conventions and inclusions are as in Supplementary Figure 14.

***Supplementary Figure 16. Control DCZ injection into OFC in the opposite hemisphere relative to the virus injection site.*** To verify whether DCZ injection has off-target effects on behavior, we conducted unilateral local injections of DCZ into the OFC, area 13m, in the left hemisphere, which is the opposite hemisphere to the virus injection site in AVMTC. The same concentration and amount of drug was injected (Methods). No significant changes were observed in overall choice probabilities, indices, or GLMs between DCZ and sham conditions, indicating no off-target effects on animal behavior. (**A-C**) – conventions are the same as in Supplementary Figures 12A, 13, and 14, respectively.

## METHODS

### Subjects

Three adult rhesus monkeys (Macaca mulatta; male, aged 8 – 10 years old) participated in the experiments. All procedures conformed to the Guide for the Care and Use of Laboratory Animals and were approved by the Washington University Institutional Animal Care and Use Committee.

### Data acquisition

We recorded single-unit neuronal activities from the anterior ventral medial temporal cortex (AVMTC) and the orbitofrontal cortex (OFC). A plastic head holder and plastic recording chambers were anchored to the skull under general anesthesia and sterile surgical conditions. Neuronal recording chambers were placed to aim at brain regions of interest. After the monkeys recovered from surgery, they participated in behavioral, electrophysiological, and DREADD experiments.

Recording sites were determined with a 1mm spacing grid system with the aid of a magnetic resonance image (3T). This MRI-based estimation of neural recording locations was aided by custom-built software (PyElectrode; Daye et al., 2013) and histology. Single-unit recording was performed using epoxy-coated tungsten electrodes (FHC) and 32- and 64-channel linear arrays (v-probes, Plexon). MR imaging with electrodes at recording sites was used to confirm recording locations within AVMTC and OFC. Electrodes or linear arrays were inserted into the brain through a stainless-steel guide tube and advanced by an oil-driven micromanipulator (MO-97A, Narishige). Signal acquisition (including amplification and filtering) was performed using an OmniPlex 40kHz recording system (Plexon). Spike sorting was performed offline using an automated spike sorting software (Kilosort2) to extract clusters from the recording data. Extracted clusters were then manually curated to identify sets of clusters corresponding to single action potentials and the periods when they were well isolated.

Among subdivisions of OFC, we mainly focused on area 13m, which is reported to play an important role in value computation and decision-making (FitzGerald et al., 2009; Padoa-Schioppa, 2007; Padoa-Schioppa & Cai, 2011). To identify area 13m, we used anatomical landmarks such as the caudate nucleus, medial orbital sulcus, and lateral orbital sulcus. The anterior-posterior extent of recording sites in the area 13m was approximately +33 to +36 interaural. AVMTC included the perirhinal cortex and TEav (medial inferior temporal cortex). To identify AVMTC, we used anatomical landmarks such as the putamen, claustrum, insula cortex, and amygdala. We also identified AVMTC as the location where strong novelty-related signals were found (Li et al., 1993; Ogasawara et al., 2022; Tamura et al., 2017; Xiang & Brown, 1998; Zhang et al., 2022; Zhu et al., 1995; Zhu & Brown, 1995). Neurons were recorded anterior-posterior extent from approximately +15 to +21 mm interaural. Example recording sites in the OFC and AVMTC are shown in Figure 4B-right and Supplementary Figure 11F.

All neuronal and behavioral analyses were conducted using MATLAB (Mathworks). Eye position was monitored with an infrared eye-tracking camera (Eyelink, SR Research). MATLAB controlled all behavioral task events and visual stimuli with Psychophysics Toolbox extensions. The liquid reward was delivered utilizing a solenoid delivery reward system (CRIST Instruments).

### Tasks

The monkeys were trained on the task shown in Figure 1. There were two contexts where offers were associated with different predictions of reward and novelty (blocks 1-2) and two other contexts served as controls where offers were associated with different predictions of reward but not novelty (block 3) and where offers were associated with different predictions of novelty but not different probabilities of reward (block 4). Each block contained five distinct offers. In block 1, five offers predicted future novel versus familiar objects with five distinct probabilities (% of novel/familiar objects: 100/0, 75/25, 50/50, 25/75, 0/100). Novel objects were associated with future big rewards, whereas familiar objects were associated with future small rewards (“novel → good”). Conversely, in block 2, five offers predicted future novel versus familiar objects with the same probabilities as block 1, but here, novel objects predicted small rewards, whereas familiar objects predicted big rewards (“novel → bad”). In block 3, five offers predicted two sets of familiar objects with five probabilities (“no novelty”). A set of familiar objects was associated with big rewards, whereas another set was associated with small rewards. In block 4, five offers predicted future novel versus familiar objects with five probabilities, but the reward size was always medium, irrespective of probabilities of novel/familiar objects (“novel → neutral”).

The task began with a fixation dot on the center of a computer screen, and the animal was required to gaze at it (Figure 1B). After 0.4s of continuous fixation, the dot was replaced by an offer. The offer remained on the screen for 1s and was then replaced by a feedback cue (a novel or familiar fractal object). Novel fractals were generated on a trial-by-trial basis, ensuring the animal had not seen them in the past (Ogasawara et al., 2022). After 1s, the feedback cue disappeared, and a juice reward was delivered as an outcome. All visual stimuli were presented in the center of the screen, and the animal was required to keep fixation on them until it received a juice reward. The fixation dot colors indicated the current block (Figure 1B). Each block consisted of 40 trials, and one of five predictions was randomly chosen for each trial. The order of the blocks was random. To ensure our key results from neural recordings were not derived from the range adaptation of neuronal activities depending on the blocked task structure (Conen & Padoa-Schioppa, 2019), monkeys also participated in an “unblocked” task where the offer could be any of the 20 offers in Figure 1A. The fixation spot color in this version of the task was held constant.

After training, monkeys also participated in a choice task (Figure 4A), where they chose among two offers drawn from the 20 offers in Figure 1A. The animal chose either of the offers by gazing at it for 0.65s. Then, the chosen offer was flipped into a feedback fractal, and the unchosen offer simultaneously disappeared. The feedback fractal remained on the screen for 2s, and the scheduled juice reward associated with the feedback cue was delivered.

### Data analyses

All statistical tests were nonparametric and two-tailed unless stated otherwise. To quantify the impact of expected reward and the interaction between expected novelty and reward on animal behavior, we created Reward and Novelty-Reward interaction indices. We used the trial-by-trial mean of lick counts 500 ms before feedback onset until feedback. The Reward index was obtained by subtracting the mean lick counts in trials with low E[R] (0% big reward, *b* in Figure 1D-left) from those with high E[R] (100% big reward, *a* in Figure 1D-left) from blocks 1-3, such that effects of novelty were counterbalanced. The Novelty-Reward interaction index was defined as the difference in the impact of expected novelty between high versus low E[R] offers, as shown in Figure 1D-right where *a*, *b*, *c*, and *d* were: the mean lick counts in trials with 100% big reward and 100% novelty from block 1; those with 100% big reward and 0% novelty from blocks 2 and 3; those with 0 % big reward and 100% novelty from block 2; and those with 0% big reward and 0% novelty from blocks 1 and 3, respectively. We calculated the same indices using choice preference (% choice). In addition, we also calculated a Novelty index to quantify the impact of expected novelty on choice (Supplementary Figure 5B). The Novelty index was analogous to the Reward index. Its calculation used the same equation as the Reward index (Figure 1D-left), but here *a* and *b* were overall choice probabilities of offers with 100 % and 0% novelty, respectively, from block 4 where reward was constant.

Each animal’s binary choice data was fit with a standard logistic regression model of choice behavior, as previously described (Bromberg-Martin et al., 2024). In effect, the log odds of choosing offer 2 over offer 1 was modeled as the difference in the values of the two offers:

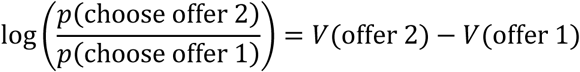

The value of each offer *i* was modeled as a linear weighted combination of the offer’s vector of *n* attributes <*x*_*i*,1_, *x*_*i*,2_, …, *x*_*i*,*n*_>:

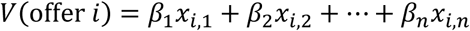

Thus, the resulting model was a GLM for binomial data with a logistic link function, with the equation:

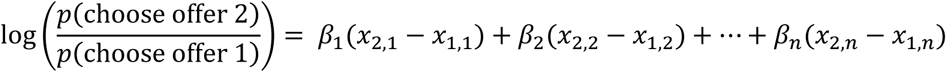

The attributes used for regression were: the main effects of expected reward (expectation of big reward) and expected novelty (expectation of novelty), and the interaction between expected reward and expected novelty. When the model was applied to choice data of the unilateral pathway manipulation (Figure 4), we included additional regressors: the main effect of the visual hemifield of chosen offers (*Contra*, 0 or 1 if the chosen offer was on ipsilateral or contralateral side), and the pairwise interactions between *Contra* and each of the following variables: expected reward, expected novelty, and the interaction between expected reward and expected novelty. In a subset of control analyses, we included a control “block effect” regressor (Supplemental Figure 6, 7, and 15). This was to confirm that our results were not affected by a behavior we noticed in some animals that preferred offers with a chance of big reward from blocks 1 and 2 (where the presence or absence of novelty predicted reward) over the corresponding offers with the same expected reward from block 3 (where novelty was absent) (Supplemental Figure 2-inset). To model this, the block effect regressor was 1 for offers from blocks 1 and 2 with a chance of big reward and 0 otherwise. Finally, when the model was applied to DREADD data, we included a binary indicator regressor that was 1 for data from the DCZ condition and 0 for data from the sham condition. To obtain regression coefficients in standardized units, all attributes were standardized by z-scoring before being used to construct the regressors (except the DREADD indicator regressor).

### Neural analyses

All single units in regions of interest were included in the analyses. In total, we obtained 837 neurons in the blocked task: 180, 27, and 78 neurons were from AVMTC in Monkeys Z, A, and L, respectively, and 147, 155, and 250 neurons were from OFC in monkeys Z, A, and L, respectively. We also obtained 698 neurons in the unblocked control version of the task: 86 in AVMTC in Monkey L, and 206 and 406 neurons were from OFC in monkeys A and L, respectively. Neurons were not excluded based on responsiveness or task variance. All analyses of offer-related neural processing were performed in the time window of 150 milliseconds after the offer presentation until the time of the feedback cue presentation (1 second after offer presentation). The window was chosen such that they included key neuronal modulations in each area and were wide enough to avoid bias towards a particular response pattern (e.g., phasic versus tonic). Neural activity was converted to normalized activity as follows. Each neuron’s spiking activity was smoothed with a Gaussian kernel (100 ms) and then z-scored. To z-score the activity, the mean and standard deviation were obtained from the neuron’s activity time course across all task conditions utilized (from 1 second before trial start fixation to 1 second after the outcome).

Across all experiments, to derive neural selectivity indices that measured the strength of discrimination among task events (e.g., Figure 2A), we used ROC area (area under curve, AUC) to distinguish spike counts across groups of trials (e.g., large reward versus small reward, novel versus familiar). For the Reward indices for blocks 1, 2, and 3, we compared trials with 100% big reward to those with 0% big reward in each block, respectively. To assess the modulation of big-reward expectation signals by novelty (Novelty-Reward index), we compared trials with 100% big reward and 100% novelty in block 1 to those with 100% big reward and 0% novelty in block 2. Significance of the ROC-based indices was assessed with Wilcoxon rank sum tests (p<0.05). When GLMs were used to analyze neural activity, we used the exact same models as previously described for behavioral analyses, and which model was used is specified in the Main text. GLM analysis of behavior provided the subjective value (weights; log odds on choice) of each offer, and these values were used in the analysis relating neural activation offer values in Figure 2B.

Novelty and reward signal latencies were derived as in other studies (Monosov et al., 2008, 2010; Ogasawara et al., 2022; White et al., 2019). These analyses were meant to assess the structure of neural activations in our task across different brain areas. Briefly, for two groups of trials (novelty-predicting and familiarity-predicting offer trials in block 4 for novelty latency analysis or large reward- and small reward-predicting offer trials in block 3 for reward latency analysis), we computed AUC in time, comparing spike density functions across the two conditions. Latency of discrimination was when the AUC was *significantly* greater or smaller than chance (0.5 AUC being chance; threshold: >=0.51 or =<0.49; p-value threshold: 0.05) for at least 50 milliseconds, continuously. We performed these analyses including all neurons in each brain area without preselection. All neurons that yielded a latency were therefore included. We also performed analysis of latency using a running GLM (same models as used to quantify behavior) moving in 1 millisecond steps and obtained the times when E[R] and E[N] were significantly modulated (P < 0.05) for at least 50 milliseconds (same as for the ROC based analyses; significant betas had to be greater than 0.05 or smaller than 0.05). Comparing latencies across areas using this analysis yielded similar results shown in the Supplementary Materials. To determine whether there was a relative latency difference across brain areas when signals were statistically observable, statistical comparisons were made across the 35% of the fastest latencies in each area using Wilcoxon rank sum tests.

### Viral vector injections

An AAV2.1 (AAV2.1-hSynI-Flag-hM4D-IRES2-AcGFP, 2 x 10^13^ genome copies per ml) vector was provided by Innovative brain virus vector core (Takada Group, Kyoto University, Japan). We unilaterally injected the viral vector into the right AVMTC of monkeys L and Z. 4 injection sites were determined where novelty-related neurons were enriched in single-unit recording (an example site is shown in Supplementary Figure 11F). The viral vector was pressure-injected, 0.1 µl per minute at 2-minute intervals (2 – 2.5 µl for each site), using a 10-µl microsyringe (Hamilton) with a handmade-injectrode. After the injection was completed, the injectrode remained in situ for 60 minutes to minimize upward leakage along the injectrode track.

### Drug injection

DCZ (Bio-Techne) was dissolved in dimethyl sulfoxide (Bio-Techne) and stored at −80 °C, shaded. This stock solution was then diluted in phosphate-buffered saline (PBS) for a final concentration of DCZ (100–300 nM for monkey L, 100–200 nM for monkey Z) on the day of use. The concentration was determined based on a previous study (Oyama et al., 2021). This solution was unilaterally injected into the OFC in the right hemisphere where AVMTC expressed hM4Di. The solution was also unilaterally injected as a control in the OFC of the opposite (left) hemisphere. DCZ injection into the opposite OFC did not impact the animal’s behavior, suggesting that our drug concentration or injecting solution itself did not have an “off-target” effect (Supplementary Figure 16). Injection sites were chosen where value-related neurons were enriched in single-unit recording (an example site is shown in Figure 4B). The drug solution was pressure-injected, 0.1 µl per 30s at 30-second intervals (5.4–8.2 µl for monkey L, 5 µl for monkey Z), using the same system as virus injection. 50 minutes after the infusion was completed, the monkeys started performing the choice task for a maximum of 3 hours (Figure 4C). As a sham experiment, we conducted precisely the same procedures as those in the DCZ injection sessions except placing the injectrode into the brain. DCZ and sham sessions are intermingled, and at least 4 days were spaced from a DCZ session to the next DCZ or sham session. All DCZ injection experiments in the same hemisphere as virus-injected AVMTC were conducted during the period when the expression level of hM4Di was high enough to effectively control neuronal activities and behavior (301–359 and 129–218 days post-virus injection for monkeys L and Z, respectively)(Nagai et al., 2024).

### Positron emission tomography (PET) imaging

PET scans were conducted at 344- and 379 days post-injection for monkeys Z and A, respectively. The animal was intubated and anesthetized for the PET scan using general anesthesia and mechanical ventilation. Isoflurane (1 – 2%) O2 (100%) was titrated to keep vitals stable for the duration of the procedure. The animal was positioned in the PET/CT scanner, and scans were acquired as outlined below.

PET/CT scans were acquired using the MRS PET/CT 220 scanner (MR Solutions Ltd, Surrey, UK). Reconstructed resolution is 1.2 mm full width half max in the center of the field of view using 3-dimensional ordered subsets expectation-maximization with 4 iterations and 16 subsets. Following scout CT image acquisition, animal head position was centered in the CT field of view, and CT attenuation images were acquired. The animal was then advanced to the center of the field of view for PET. [^11^C]DCZ (525 MBq for monkey L, 394 MBq for monkey Z) was intravenously injected over 20 seconds through the saphenous vein, followed by a 5 ml 0.9% sodium chloride flush. PET data acquisition began simultaneously with the initiation of radioisotope administration, and dynamic PET images were collected for 120 minutes. Images were corrected for attenuation, scatter, randoms, and dead time and then binned over 24 frames (3×60s, 4×120s, 3×180s, 8×300s, 6×600s).

### PET data processing

The early frames were combined to obtain epochs of duration 4 – 10 min to ensure adequate counting statistics for purposes of image registration (Snyder, 1996). The resulting 18 time-sliced images were spatially smoothed (4mm Gaussian pre-blur in each direction) and mutually co-registered (6-parameter rigid body transform) in a manner that minimizes the discrepancy between all transforms and their inverses representing all image pairs analyzed as a set (Eisenstein et al., 2012). The pre-blur parameter value (4mm) was previously optimized as that which minimizes the global transform discrepancy (optimized value = 0.09 rms mm, 0.26 rms deg). This procedure yielded 18 transforms compensating for head motion (< 1 mm translation, < 1 deg rotation) occurring during the [^11^C]DCZ scan.

An average [^11^C]DCZ PET emission image was generated using all time slices. This image then was registered (6-parameter rigid body transform, the multimodal objective function; (Rowland et al., 2005)) to a T1-weighted (T1w) structural MRI (MPRAGE) previously acquired. The T1w image was registered (12-parameter affine transform) to our laboratory-specific atlas-representative T1w target image. Finally, composition of transforms (time sliced PET image → PET data average → T1w → atlas) enabled resampling all 18 unblurred, time sliced [^11^C]DCZ images in atlas space.

Specific [^11^C]DCZ uptake was evaluated in an unblurred image summed over post-injection minutes 90-120. A region of interest (ROI) representing [^11^C]DCZ-specific uptake was generated by clustering and thresholding the image shown in Supplementary Figure 11A-B. Additional ROIs were defined as 2.5mm radius spheres, visually centered on the homologous (L) perirhinal region, the R putamen, and R subcortical white matter. Time-activity curves (Supplementary Figure 11 C-D) were generated by evaluating the time-sliced data within all ROIs.

### Histology

After the end of the DREADD experiment, the monkey was deeply anesthetized with sodium pentobarbital and transcardially perfused with phosphate-buffered saline (PBS), followed by 4% paraformaldehyde (PFA) in PBS (pH 7.4). The brain was postfixed in 4% PFA overnight, blocked, and equilibrated with 30% sucrose in PBS at 4 °C. Frozen sections were then cut on a sliding microtome at 50µm in the coronal plane.

To visualize the immunoreactive signals of green fluorescent protein (GFP) co-expressed with hM4Di, we followed the methodology described in a study (Oyama et al., 2021). Sections of ROIs were immersed in 1% skim milk containing 0.1% TritonX-100 for 1 hour at room temperature and washed 1x with blocking buffer (1x PBS containing 0.1% TX-100 and 1% normal goat serum) for 15 minutes at room temperature. The sections were then incubated overnight at 4 °C with rabbit anti-GFP monoclonal antibody (1:500; G10362, Thermo Fisher Scientific) in blocking buffer for 2 days at 4 °C. The sections were washed 3×10 minutes with 1x PBS and then incubated in the same fresh medium containing biotinylated goat anti-rabbit immunoglobulin G antibody (1:1000; Jackson ImmunoResearch) for 2 hours at room temperature, washed 3×10 minutes with 1x PBS, incubated in avidin-biotin-peroxidase complex (ABC Elite, Vector Laboratories) for 2 hours at room temperature, and washed another 3×10 minutes with 1x PBS.

To visualize the antigen, the sections were reacted in 3,3-diaminobenzidine (DAB) with cobalt chloride and nickel chloride metal enhancers (Thermo Fisher 34065). The sections were mounted on gelatin-coated glass slides, air-dried, dehydrated in an ethanol series, cleared with xylene, and cover-slipped with DPX (#06522, Sigma). The sections were then imaged under 10 and 60x resolutions for the whole brain slice and axon boutons, respectively.

## Acknowledgements

This work was supported by the National Institute of Mental Health under award numbers R01MH128344, R01MH110594, and R01MH116937, and Conte Center on the Neurocircuitry of OCD MH10643, and Paula & Rodger Riney Foundation (to J.S.P), and Barnes-Jewish Hospital Foundation (to J.S.P). The viral vector is supplied with the supports of the program for Brain Mapping by Integrated Neurotechnologies for Disease Studies (Brain/MINDS) from the Japan Agency for Medical Research and Development (AMED) under grant number JP24wm0625103 (to K.I.), and of the Japan Society for the Promotion of Science under grant number JP22H05157 (to K.I.). We are grateful to Kim Kocher for the animal care and animal training, to Andreas Burkhalter and Rebecca Mellor for their histology services and scientific advice, to Peter Bayguinov for performing tissue imaging, to Aaron Tanenbaum for creating the T1w atlas used in PET analysis, to Ethan S. Bromberg-Martin for giving us valuable comments on the various aspects of the analyses, Kei Kimura for advice on tissue staining, and to Kei Oyama for advice on GFP visualization.

## Author contributions

TO and IEM conceptualized the research. TO performed the experiments and analyzed the data. KX assisted TO in data acquisition and in other aspects of the experiments. JSP supervised the PET imaging. AZS analyzed the PET data and advised the team on imaging-data interpretation. ZT provided and synthesized the radioactive agent utilized in the PET imaging. TM, KI, and MT provided the viral vector and provided scientific advice on its utilization. TO and IEM wrote the manuscript. IEM assisted with analyses, guided the research, and acquired the funding.

## REFERENCES

Agster, K. L., & Burwell, R. D. (2009). Cortical efferents of the perirhinal, postrhinal, and entorhinal cortices of the rat. Hippocampus, 19(12), 1159–1186. 10.1002/hipo.20578

Ahmadlou, M., Houba, J. H. W., Vierbergen, J. F. M. V., Giannouli, M., Gimenez, G.-A., Weeghel, C. V., Darbanfouladi, M., Shirazi, M. Y., Dziubek, J., Kacem, M., Winter, F. D., & Heimel, J. A. (2021). A cell type–specific cortico-subcortical brain circuit for investigatory and novelty-seeking behavior. Science, 372(6543), eabe9681. 10.1126/science.abe9681

Aubret, A., Matignon, L., & Hassas, S. (2023). An Information-Theoretic Perspective on Intrinsic Motivation in Reinforcement Learning: A Survey. Entropy, 25(2), Article 2. 10.3390/e25020327

Ballesta, S., Shi, W., Conen, K. E., & Padoa-Schioppa, C. (2020). Values encoded in orbitofrontal cortex are causally related to economic choices. Nature, 588(7838), 450–453. 10.1038/s41586-020-2880-x

Barbas, H. (1993). Organization of cortical afferent input to orbitofrontal areas in the rhesus monkey. Neuroscience, 56(4), 841–864. 10.1016/0306-4522(93)90132-Y

Bellemare, M. G., Srinivasan, S., Ostrovski, G., Schaul, T., Saxton, D., & Munos, R. (2016). Unifying Count-Based Exploration and Intrinsic Motivation. Advances in Neural Information Processing Systems, 29. https://proceedings.neurips.cc/paper/2016/hash/afda332245e2af431fb7b672a68b659d-Abstract.html

Berlyne, D. E. (1950). Novelty and curiosity as determinants of exploratory behaviour. British Journal of Psychology. General Section, 41(1), 68–81.

Berlyne, D. E. (1954). A Theory of Human Curiosity. British Journal of Psychology. General Section, 45(3), 180–191. 10.1111/j.2044-8295.1954.tb01243.x

Berlyne, D. E. (1966). Curiosity and Exploration. Science, 153(3731), 25–33. 10.1126/science.153.3731.25

Bromberg-Martin, E. S., Feng, Y.-Y., Ogasawara, T., White, J. K., Zhang, K., & Monosov, I. E. (2024). A neural mechanism for conserved value computations integrating information and rewards. Nature Neuroscience, 27(1), 159–175. 10.1038/s41593-023-01511-4

Brown, M. W., & Aggleton, J. P. (2001). Recognition memory: What are the roles of the perirhinal cortex and hippocampus? Nature Reviews Neuroscience, 2(1), 51–61. 10.1038/35049064

Burda, Y., Edwards, H., Storkey, A., & Klimov, O. (2018). Exploration by Random Network Distillation (No. arXiv:1810.12894). *arXiv*. 10.48550/arXiv.1810.12894

Burns, L. H., Annett, L., Kelley, A. E., Everitt, B. J., & Robbins, T. W. (1996). Effects of lesions to amygdala, ventral subiculum, medial prefrontal cortex, and nucleus accumbens on the reaction to novelty: Implication for limbic—striatal interactions. Behavioral Neuroscience, 110(1), 60–73. 10.1037/0735-7044.110.1.60

Bussey, T. J., & Saksida, L. M. (2002). The organization of visual object representations: A connectionist model of effects of lesions in perirhinal cortex. European Journal of Neuroscience, 15(2), 355–364. 10.1046/j.0953-816x.2001.01850.x

Bussey, T. J., Saksida, L. M., & Murray, E. A. (2005). The Perceptual-Mnemonic/Feature Conjunction Model of Perirhinal Cortex Function. The Quarterly Journal of Experimental Psychology Section B, 58(3–4b), 269–282. 10.1080/02724990544000004

Caplin, A., & Dean, M. (2015). Revealed Preference, Rational Inattention, and Costly Information Acquisition. American Economic Review, 105(7), 2183–2203. 10.1257/aer.20140117

Caplin, A., & Leahy, J. (2001). Psychological Expected Utility Theory and Anticipatory Feelings*. The Quarterly Journal of Economics, 116(1), 55–79. 10.1162/003355301556347

Carmichael, S. T., & Price, J. L. (1995). Limbic connections of the orbital and medial prefrontal cortex in macaque monkeys. Journal of Comparative Neurology, 363(4), 615–641. 10.1002/cne.903630408

Chen, S., He, L., Huang, A. J. Y., Boehringer, R., Robert, V., Wintzer, M. E., Polygalov, D., Weitemier, A. Z., Tao, Y., Gu, M., Middleton, S. J., Namiki, K., Hama, H., Therreau, L., Chevaleyre, V., Hioki, H., Miyawaki, A., Piskorowski, R. A., & McHugh, T. J. (2020). A hypothalamic novelty signal modulates hippocampal memory. Nature, 586(7828), 270–274. 10.1038/s41586-020-2771-1

Cichon, J., & Gan, W.-B. (2015). Branch-specific dendritic Ca2+ spikes cause persistent synaptic plasticity. Nature, 520(7546), 180–185. 10.1038/nature14251

Clark, A. M., Bouret, S., Young, A. M., Murray, E. A., & Richmond, B. J. (2013). Interaction Between Orbital Prefrontal and Rhinal Cortex Is Required for Normal Estimates of Expected Value. Journal of Neuroscience, 33(5), 1833–1845. 10.1523/JNEUROSCI.3605-12.2013

Cohen, J. D., McClure, S. M., & Yu, A. J. (2007). Should I stay or should I go? How the human brain manages the trade-off between exploitation and exploration. Philosophical Transactions of the Royal Society B: Biological Sciences, 362(1481), 933–942. 10.1098/rstb.2007.2098

Conen, K. E., & Padoa-Schioppa, C. (2019). Partial Adaptation to the Value Range in the Macaque Orbitofrontal Cortex. Journal of Neuroscience, 39(18), 3498–3513. 10.1523/JNEUROSCI.2279-18.2019

Costa, V. D., Mitz, A. R., & Averbeck, B. B. (2019). Subcortical Substrates of Explore-Exploit Decisions in Primates. Neuron, 103(3), 533–545.e5. 10.1016/j.neuron.2019.05.017

Costa, V. D., Tran, V. L., Turchi, J., & Averbeck, B. B. (2014). Dopamine modulates novelty seeking behavior during decision making. Behavioral Neuroscience, 128(5), 556–566. 10.1037/a0037128

Daye, P. M., Monosov, I. E., Hikosaka, O., Leopold, D. A., & Optican, L. M. (2013). pyElectrode: An open-source tool using structural MRI for electrode positioning and neuron mapping. Journal of Neuroscience Methods, 213(1), 123–131. 10.1016/j.jneumeth.2012.12.012

Djamshidian, A., O’Sullivan, S. S., Wittmann, B. C., Lees, A. J., & Averbeck, B. B. (2011). Novelty seeking behaviour in Parkinson’s disease. Neuropsychologia, 49(9), 2483–2488. 10.1016/j.neuropsychologia.2011.04.026

Donfrancesco, R., Di Trani, M., Porfirio, M. C., Giana, G., Miano, S., & Andriola, E. (2015). Might the temperament be a bias in clinical study on attention-deficit hyperactivity disorder (ADHD)?: Novelty Seeking dimension as a core feature of ADHD. Psychiatry Research, 227(2), 333–338. 10.1016/j.psychres.2015.02.014

Doron, G., Shin, J. N., Takahashi, N., Drüke, M., Bocklisch, C., Skenderi, S., De Mont, L., Toumazou, M., Ledderose, J., Brecht, M., Naud, R., & Larkum, M. E. (2020). Perirhinal input to neocortical layer 1 controls learning. Science, 370(6523), eaaz3136. 10.1126/science.aaz3136

Dubey, R., & Griffiths, T. L. (2020). Reconciling novelty and complexity through a rational analysis of curiosity. Psychological Review, 127(3), 455–476. 10.1037/rev0000175

Duuren, E. van, Lankelma, J., & Pennartz, C. M. A. (2008). Population Coding of Reward Magnitude in the Orbitofrontal Cortex of the Rat. Journal of Neuroscience, 28(34), 8590–8603. 10.1523/JNEUROSCI.5549-07.2008

Duuren, E. van, Plasse, G. van der, Lankelma, J., Joosten, R. N. J. M. A., Feenstra, M. G. P., & Pennartz, C. M. A. (2009). Single-Cell and Population Coding of Expected Reward Probability in the Orbitofrontal Cortex of the Rat. Journal of Neuroscience, 29(28), 8965– 8976. 10.1523/JNEUROSCI.0005-09.2009

Easton, A., & Gaffan, D. (2000). Comparison of perirhinal cortex ablation and crossed unilateral lesions of the medial forebrain bundle from the inferior temporal cortex in the rhesus monkey: Effects on learning and retrieval. Behavioral Neuroscience, 114(6), 1041–1057. 10.1037/0735-7044.114.6.1041

Eisenstein, S. A., Koller, J. M., Piccirillo, M., Kim, A., Antenor-Dorsey, J. A. V., Videen, T. O., Snyder, A. Z., Karimi, M., Moerlein, S. M., Black, K. J., Perlmutter, J. S., & Hershey, T. (2012). Characterization of extrastriatal D2 in vivo specific binding of [18F](N-methyl)benperidol using PET. Synapse, 66(9), 770–780. 10.1002/syn.21566

Eldridge, M. A. G., Lerchner, W., Saunders, R. C., Kaneko, H., Krausz, K. W., Gonzalez, F. J., Ji, B., Higuchi, M., Minamimoto, T., & Richmond, B. J. (2016). Chemogenetic disconnection of monkey orbitofrontal and rhinal cortex reversibly disrupts reward value. Nature Neuroscience, 19(1), 37–39. 10.1038/nn.4192

Eldridge, M. A., Hines, B. E., & Murray, E. A. (2021). The visual prefrontal cortex of anthropoids: Interaction with temporal cortex in decision making and its role in the making of ‘visual animals.’ Current Opinion in Behavioral Sciences, 41, 22–29. 10.1016/j.cobeha.2021.02.012

Eldridge, M. A., Matsumoto, N., Wittig, J. H., Jnr, Masseau, E. C., Saunders, R. C., & Richmond, B. J. (2018). Perceptual processing in the ventral visual stream requires area TE but not rhinal cortex. eLife, 7, e36310. 10.7554/eLife.36310

Elston, T. W., & Wallis, J. D. (2025). Context-dependent decision-making in the primate hippocampal–prefrontal circuit. Nature Neuroscience, 28(2), 374–382. 10.1038/s41593-024-01839-5

Feierstein, C. E., Quirk, M. C., Uchida, N., Sosulski, D. L., & Mainen, Z. F. (2006). Representation of Spatial Goals in Rat Orbitofrontal Cortex. Neuron, 51(4), 495–507. 10.1016/j.neuron.2006.06.032

FitzGerald, T. H. B., Seymour, B., & Dolan, R. J. (2009). The Role of Human Orbitofrontal Cortex in Value Comparison for Incommensurable Objects. The Journal of Neuroscience, 29(26), 8388–8395. 10.1523/JNEUROSCI.0717-09.2009

Freedman, D. J., Riesenhuber, M., Poggio, T., & Miller, E. K. (2001). Categorical Representation of Visual Stimuli in the Primate Prefrontal Cortex. Science, 291(5502), 312–316. 10.1126/science.291.5502.312

Freedman, D. J., Riesenhuber, M., Poggio, T., & Miller, E. K. (2003). A Comparison of Primate Prefrontal and Inferior Temporal Cortices during Visual Categorization. The Journal of Neuroscience, 23(12), 5235–5246. 10.1523/JNEUROSCI.23-12-05235.2003

Gittins, J., Glazebrook, K., & Weber, R. (2011). Multi-armed Bandit Allocation Indices. John Wiley & Sons.

Gottlieb, J., Lopes, M., & Oudeyer, P.-Y. (2016). Motivated Cognition: Neural and Computational Mechanisms of Curiosity, Attention, and Intrinsic Motivation. In S. Kim, J. Reeve, & M. Bong (Eds.), Advances in Motivation and Achievement (Vol. 19, pp. 149–172). Emerald Group Publishing Limited. 10.1108/S0749-742320160000019017

Gottlieb, J., & Oudeyer, P.-Y. (2018). Towards a neuroscience of active sampling and curiosity. Nature Reviews Neuroscience, 19(12), 758–770. 10.1038/s41583-018-0078-0

Hall, G. S., & Smith, T. L. (1903). Curiosity and Interest. The Pedagogical Seminary, 10(3), 315–358. 10.1080/08919402.1903.10532722

Houillon, A., Lorenz, R. C., Boehmer, W., Rapp, M. A., Heinz, A., Gallinat, J., & Obermayer, K. (2013). The effect of novelty on reinforcement learning. In Progress in Brain Research (Vol. 202, pp. 415–439). Elsevier. 10.1016/B978-0-444-62604-2.00021-6

Hunt, L. T., Malalasekera, W. M. N., de Berker, A. O., Miranda, B., Farmer, S. F., Behrens, T. E. J., & Kennerley, S. W. (2018). Triple dissociation of attention and decision computations across prefrontal cortex. Nature Neuroscience, 21(10), 1471–1481. 10.1038/s41593-018-0239-5

Jaegle, A., Mehrpour, V., & Rust, N. (2019). Visual novelty, curiosity, and intrinsic reward in machine learning and the brain. Current Opinion in Neurobiology, 58, 167–174. 10.1016/j.conb.2019.08.004

James, W. (1913). The Principles of Psychology. Henry Holt.

James, W. (1983). Talks to Teachers on Psychology and to Students on Some of Life’s Ideals. Harvard University Press.

Kahneman, D., & Tversky, A. (1979). Prospect Theory: An Analysis of Decision under Risk. Econometrica, 47, 263–292.

Kakade, S., & Dayan, P. (2002). Dopamine: Generalization and bonuses. Neural Networks, 15(4–6), 549–559. 10.1016/S0893-6080(02)00048-5

Kim, S. W., & Grant, J. E. (2001). Personality dimensions in pathological gambling disorder and obsessive–compulsive disorder. Psychiatry Research, 104(3), 205–212. 10.1016/S0165-1781(01)00327-4

Kim, Y., Nam, W., Kim, H., Kim, J.-H., & Kim, G. (2019). Curiosity-Bottleneck: Exploration by Distilling Task-Specific Novelty. Proceedings of the 36th International Conference on Machine Learning, 3379–3388. https://proceedings.mlr.press/v97/kim19c.html

Kimura, K., Nagai, Y., Hatanaka, G., Fang, Y., Tanabe, S., Zheng, A., Fujiwara, M., Nakano, M., Hori, Y., Takeuchi, R. F., Inagaki, M., Minamimoto, T., Fujita, I., Inoue, K., & Takada, M. (2023). A mosaic adeno-associated virus vector as a versatile tool that exhibits high levels of transgene expression and neuron specificity in primate brain. Nature Communications, 14(1), 4762. 10.1038/s41467-023-40436-1

Knight, F. H. (1921). Risk, Uncertainty and Profit. Houghton Mifflin.

Knudsen, E. B., & Wallis, J. D. (2022). Taking stock of value in the orbitofrontal cortex. Nature Reviews Neuroscience, 23(7), 428–438. 10.1038/s41583-022-00589-2

Kreitler, S., Zigler, E., & Kreitler, H. (1975). The nature of curiosity in children. Journal of School Psychology, 13(3), 185–200. 10.1016/0022-4405(75)90002-3

Kreps, D. M., & Porteus, E. L. (1978). Temporal Resolution of Uncertainty and Dynamic Choice Theory. Econometrica, 46(1), 185–200. 10.2307/1913656

Kusunoki, K., Sato, T., Taga, C., Yoshida, Y., Komori, K., Narita, T., Hirano, S., Iwata, N., & Ozaki, N. (2000). Low novelty-seeking differentiates obsessive-compulsive disorder from major depression. Acta Psychiatrica Scandinavica, 101(5), 403–405. 10.1034/j.1600-0447.2000.101005403.x

Lacefield, C. O., Pnevmatikakis, E. A., Paninski, L., & Bruno, R. M. (2019). Reinforcement Learning Recruits Somata and Apical Dendrites across Layers of Primary Sensory Cortex. Cell Reports, 26(8), 2000–2008.e2. 10.1016/j.celrep.2019.01.093

Lak, A., Stauffer, W. R., & Schultz, W. (2016). Dopamine neurons learn relative chosen value from probabilistic rewards. eLife, 5, e18044. 10.7554/eLife.18044

Lavenex, P., Suzuki, W. A., & Amaral, D. G. (2002). Perirhinal and parahippocampal cortices of the macaque monkey: Projections to the neocortex. Journal of Comparative Neurology, 447(4), 394–420. 10.1002/cne.10243

Leinweber, M., Ward, D. R., Sobczak, J. M., Attinger, A., & Keller, G. B. (2017). A Sensorimotor Circuit in Mouse Cortex for Visual Flow Predictions. Neuron, 95(6), 1420–1432.e5. 10.1016/j.neuron.2017.08.036

Li, L., Miller, E. K., & Desimone, R. (1993). The representation of stimulus familiarity in anterior inferior temporal cortex. Journal of Neurophysiology, 69(6), 1918–1929. 10.1152/jn.1993.69.6.1918

Lim, S.-L., O’Doherty, J. P., & Rangel, A. (2011). The Decision Value Computations in the vmPFC and Striatum Use a Relative Value Code That is Guided by Visual Attention. The Journal of Neuroscience, 31(37), 13214–13223. 10.1523/JNEUROSCI.1246-11.2011

Manita, S., Suzuki, T., Homma, C., Matsumoto, T., Odagawa, M., Yamada, K., Ota, K., Matsubara, C., Inutsuka, A., Sato, M., Ohkura, M., Yamanaka, A., Yanagawa, Y., Nakai, J., Hayashi, Y., Larkum, M. E., & Murayama, M. (2015). A Top-Down Cortical Circuit for Accurate Sensory Perception. Neuron, 86(5), 1304–1316. 10.1016/j.neuron.2015.05.006

McGinty, V. B., Rangel, A., & Newsome, W. T. (2016). Orbitofrontal Cortex Value Signals Depend on Fixation Location during Free Viewing. Neuron, 90(6), 1299–1311. 10.1016/j.neuron.2016.04.045

McKee, J. L., Riesenhuber, M., Miller, E. K., & Freedman, D. J. (2014). Task Dependence of Visual and Category Representations in Prefrontal and Inferior Temporal Cortices. Journal of Neuroscience, 34(48), 16065–16075. 10.1523/JNEUROSCI.1660-14.2014

Menegas, W., Babayan, B. M., Uchida, N., & Watabe-Uchida, M. (2017). Opposite initialization to novel cues in dopamine signaling in ventral and posterior striatum in mice. eLife, 6, e21886. 10.7554/eLife.21886

Miyashita, Y. (2019). Perirhinal circuits for memory processing. Nature Reviews Neuroscience, 20(10), 577–592. 10.1038/s41583-019-0213-6

Modirshanechi, A., Kondrakiewicz, K., Gerstner, W., & Haesler, S. (2023). Curiosity-driven exploration: Foundations in neuroscience and computational modeling. Trends in Neurosciences, 46(12), 1054–1066. 10.1016/j.tins.2023.10.002

Modirshanechi, A., Lin, W.-H., Xu, H. A., Herzog, M. H., & Gerstner, W. (2023). The curse of optimism: A persistent distraction by novelty (p. 2022.07.05.498835). *bioRxiv*. 10.1101/2022.07.05.498835

Monosov, I. E. (2024). Curiosity: Primate neural circuits for novelty and information seeking. Nature Reviews Neuroscience, 25(3), 195–208. 10.1038/s41583-023-00784-9

Monosov, I. E., Leopold, D. A., & Hikosaka, O. (2015). Neurons in the Primate Medial Basal Forebrain Signal Combined Information about Reward Uncertainty, Value, and Punishment Anticipation. Journal of Neuroscience, 35(19), 7443–7459. 10.1523/JNEUROSCI.0051-15.2015

Monosov, I. E., Ogasawara, T., Haber, S. N., Heimel, J. A., & Ahmadlou, M. (2022). The zona incerta in control of novelty seeking and investigation across species. Current Opinion in Neurobiology, 77, 102650. 10.1016/j.conb.2022.102650

Monosov, I. E., Sheinberg, D. L., & Thompson, K. G. (2010). Paired neuron recordings in the prefrontal and inferotemporal cortices reveal that spatial selection precedes object identification during visual search. Proceedings of the National Academy of Sciences, 107(29), 13105–13110. 10.1073/pnas.1002870107

Monosov, I. E., Trageser, J. C., & Thompson, K. G. (2008). Measurements of Simultaneously Recorded Spiking Activity and Local Field Potentials Suggest that Spatial Selection Emerges in the Frontal Eye Field. Neuron, 57(4), 614–625. 10.1016/j.neuron.2007.12.030

Morecraft, R. J., Geula, C., & Mesulam, M.-M. (1992). Cytoarchitecture and neural afferents of orbitofrontal cortex in the brain of the monkey. Journal of Comparative Neurology, 323(3), 341–358. 10.1002/cne.903230304

Murayama, K., FitzGibbon, L., & Sakaki, M. (2019). Process Account of Curiosity and Interest: A Reward-Learning Perspective. Educational Psychology Review, 31(4), 875–895. 10.1007/s10648-019-09499-9

Murray, E. A., & Richmond, B. J. (2001). Role of perirhinal cortex in object perception, memory, and associations. Current Opinion in Neurobiology, 11(2), 188–193. 10.1016/S0959-4388(00)00195-1

Nagai, Y., Hori, Y., Inoue, K., Hirabayashi, T., Mimura, K., Oyama, K., Miyakawa, N., Hori, Y., Iwaoki, H., Kumata, K., Zhang, M.-R., Takada, M., Higuchi, M., & Minamimoto, T. (2024). Longitudinal assessment of DREADD expression and efficacy in the monkey brain (p. 2024.12.26.630299). bioRxiv. 10.1101/2024.12.26.630299

Nagai, Y., Miyakawa, N., Takuwa, H., Hori, Y., Oyama, K., Ji, B., Takahashi, M., Huang, X.-P., Slocum, S. T., DiBerto, J. F., Xiong, Y., Urushihata, T., Hirabayashi, T., Fujimoto, A., Mimura, K., English, J. G., Liu, J., Inoue, K., Kumata, K., … Minamimoto, T. (2020). Deschloroclozapine, a potent and selective chemogenetic actuator enables rapid neuronal and behavioral modulations in mice and monkeys. Nature Neuroscience, 23(9), 1157–1167. 10.1038/s41593-020-0661-3

Ogasawara, T., Sogukpinar, F., Zhang, K., Feng, Y.-Y., Pai, J., Jezzini, A., & Monosov, I. E. (2022). A primate temporal cortex–zona incerta pathway for novelty seeking. Nature Neuroscience, 25(1), 50–60. 10.1038/s41593-021-00950-1

Oudeyer, P.-Y., Kaplan, F., & Hafner, V. V. (2007). Intrinsic Motivation Systems for Autonomous Mental Development. IEEE Transactions on Evolutionary Computation, 11(2), 265–286. 10.1109/TEVC.2006.890271

Oyama, K., Hori, Y., Nagai, Y., Miyakawa, N., Mimura, K., Hirabayashi, T., Inoue, K., Suhara, T., Takada, M., Higuchi, M., & Minamimoto, T. (2021). Chemogenetic dissection of the primate prefronto-subcortical pathways for working memory and decision-making. Science Advances, 7(26), eabg4246. 10.1126/sciadv.abg4246

Padoa-Schioppa, C. (2007). Orbitofrontal Cortex and the Computation of Economic Value. Annals of the New York Academy of Sciences, 1121(1), 232–253. 10.1196/annals.1401.011

Padoa-Schioppa, C., & Assad, J. A. (2006). Neurons in the orbitofrontal cortex encode economic value. Nature, 441(7090), 223–226. 10.1038/nature04676

Padoa-Schioppa, C., & Cai, X. (2011). The orbitofrontal cortex and the computation of subjective value: Consolidated concepts and new perspectives. Annals of the New York Academy of Sciences, 1239(1), 130–137. 10.1111/j.1749-6632.2011.06262.x

Padoa-Schioppa, C., & Conen, K. E. (2017). Orbitofrontal Cortex: A Neural Circuit for Economic Decisions. Neuron, 96(4), 736–754. 10.1016/j.neuron.2017.09.031

Parker, A., & Gaffan, D. (1998). Memory after frontal/temporal disconnection in monkeys: Conditional and non-conditional tasks, unilateral and bilateral frontal lesions. Neuropsychologia, 36(3), 259–271. 10.1016/S0028-3932(97)00112-7

Pelletier, G., Aridan, N., Fellows, L. K., & Schonberg, T. (2021). A Preferential Role for Ventromedial Prefrontal Cortex in Assessing “the Value of the Whole” in Multiattribute Object Evaluation. Journal of Neuroscience, 41(23), 5056–5068. 10.1523/JNEUROSCI.0241-21.2021

Pelletier, G., & Fellows, L. K. (2021). Viewing orbitofrontal cortex contributions to decision-making through the lens of object recognition. Behavioral Neuroscience, 135(2), 182–191. 10.1037/bne0000447

Perkins, A. Q., & Rich, E. L. (2024). Attention-dependent attribute comparisons underlie multi-attribute decision-making in orbitofrontal cortex. bioRxiv. 10.1101/2024.11.12.623291

Poli, F., O’Reilly, J. X., Mars, R. B., & Hunnius, S. (2024). Curiosity and the dynamics of optimal exploration. Trends in Cognitive Sciences, 28(5), 441–453. 10.1016/j.tics.2024.02.001

Ranganathan, G. N., Apostolides, P. F., Harnett, M. T., Xu, N.-L., Druckmann, S., & Magee, J. C. (2018). Active dendritic integration and mixed neocortical network representations during an adaptive sensing behavior. Nature Neuroscience, 21(11), 1583–1590. 10.1038/s41593-018-0254-6

Rempel-Clower, N. L., & Barbas, H. (2000). The Laminar Pattern of Connections between Prefrontal and Anterior Temporal Cortices in the Rhesus Monkey is Related to Cortical Structure and Function. Cerebral Cortex, 10(9), 851–865. 10.1093/cercor/10.9.851

Roesch, M. R., & Olson, C. R. (2004). Neuronal Activity Related to Reward Value and Motivation in Primate Frontal Cortex. Science, 304(5668), 307–310. 10.1126/science.1093223

Roitman, J. D., & Roitman, M. F. (2010). Risk-preference differentiates orbitofrontal cortex responses to freely chosen reward outcomes. European Journal of Neuroscience, 31(8), 1492–1500. 10.1111/j.1460-9568.2010.07169.x

Rolls, E. T. (2000). The Orbitofrontal Cortex and Reward. Cerebral Cortex, 10(3), 284–294. 10.1093/cercor/10.3.284

Rolls, E. T., & Baylis, L. L. (1994). Gustatory, olfactory, and visual convergence within the primate orbitofrontal cortex. Journal of Neuroscience, 14(9), 5437–5452. 10.1523/JNEUROSCI.14-09-05437.1994

Rowland, D. J., Garbow, J. R., Laforest, R., & Snyder, A. Z. (2005). Registration of [18F]FDG microPET and small-animal MRI. Nuclear Medicine and Biology, 32(6), 567–572. 10.1016/j.nucmedbio.2005.05.002

Rushworth, M. F. S., Noonan, M. P., Boorman, E. D., Walton, M. E., & Behrens, T. E. (2011). Frontal Cortex and Reward-Guided Learning and Decision-Making. Neuron, 70(6), 1054–1069. 10.1016/j.neuron.2011.05.014

Sasikumar, D., Emeric, E., Stuphorn, V., & Connor, C. E. (2018). First-Pass Processing of Value Cues in the Ventral Visual Pathway. Current Biology, 28(4), 538–548.e3. 10.1016/j.cub.2018.01.051

Savinov, N., Raichuk, A., Marinier, R., Vincent, D., Pollefeys, M., Lillicrap, T., & Gelly, S. (2019). Episodic Curiosity through Reachability (No. arXiv:1810.02274). *arXiv*. 10.48550/arXiv.1810.02274

Schoenbaum, G., Chiba, A. A., & Gallagher, M. (1998). Orbitofrontal cortex and basolateral amygdala encode expected outcomes during learning. Nature Neuroscience, 1(2), 155–159. 10.1038/407

Schoenbaum, G., & Eichenbaum, H. (1995). Information coding in the rodent prefrontal cortex. I. Single-neuron activity in orbitofrontal cortex compared with that in pyriform cortex. Journal of Neurophysiology, 74(2), 733–750. 10.1152/jn.1995.74.2.733

Smock, C. D., & Holt, B. G. (1962). Children’s Reactions to Novelty: An Experimental Study of “Curiosity Motivation.” Child Development, 33(3), 631–642. 10.2307/1126663

Snyder, A. Z. (1996). Difference Image vs Ratio Image Error Function Forms in PET—PET Realignment. Quantification of Brain Function Using PET, 131–137. 10.1016/b978-012389760-2/50028-1

Sul, J. H., Kim, H., Huh, N., Lee, D., & Jung, M. W. (2010). Distinct Roles of Rodent Orbitofrontal and Medial Prefrontal Cortex in Decision Making. Neuron, 66(3), 449–460. 10.1016/j.neuron.2010.03.033

Suzuki, W. A., & Amaral, D. G. (1994). Perirhinal and parahippocampal cortices of the macaque monkey: Cortical afferents. Journal of Comparative Neurology, 350(4), 497–533. 10.1002/cne.903500402

Suzuki, W. A., & Naya, Y. (2014). The Perirhinal Cortex. Annual Review of Neuroscience, 37(1), 39–53. 10.1146/annurev-neuro-071013-014207

Takahashi, N., Ebner, C., Sigl-Glöckner, J., Moberg, S., Nierwetberg, S., & Larkum, M. E. (2020). Active dendritic currents gate descending cortical outputs in perception. Nature Neuroscience, 23(10), 1277–1285. 10.1038/s41593-020-0677-8

Takahashi, N., Oertner, T. G., Hegemann, P., & Larkum, M. E. (2016). Active cortical dendrites modulate perception. Science, 354(6319), 1587–1590. 10.1126/science.aah6066

Tamura, K., Takeda, M., Setsuie, R., Tsubota, T., Hirabayashi, T., Miyamoto, K., & Miyashita, Y. (2017). Conversion of object identity to object-general semantic value in the primate temporal cortex. Science, 357(6352), 687–692. 10.1126/science.aan4800

Tang, H., Houthooft, R., Foote, D., Stooke, A., Chen, X., Duan, Y., Schulman, J., DeTurck, F., & Abbeel, P. (2017). #Exploration: A Study of Count-Based Exploration for Deep Reinforcement Learning. Advances in Neural Information Processing Systems, 30. https://proceedings.neurips.cc/paper_files/paper/2017/hash/3a20f62a0af1aa152670bab3 c602feed-Abstract.html

Tapper, A. R., & Molas, S. (2020). Midbrain circuits of novelty processing. Neurobiology of Learning and Memory, 176, 107323. 10.1016/j.nlm.2020.107323

Thornton, J. A., Malkova, L., & Murray, E. A. (1998). Rhinal cortex ablations fail to disrupt reinforcer devaluation effects in rhesus monkeys (Macaca mulatta). Behavioral Neuroscience, 112(4), 1020–1025. 10.1037/0735-7044.112.4.1020

Thorpe, S. J., Rolls, E. T., & Maddison, S. (1983). The orbitofrontal cortex: Neuronal activity in the behaving monkey. Experimental Brain Research, 49(1), 93–115. 10.1007/BF00235545

Tian, L., Xia, Y., Flores, H. P., Campbell, M. C., Moerlein, S. M., & Perlmutter, J. S. (2015). Neuroimaging Analysis of the Dopamine Basis for Apathetic Behaviors in an MPTP-Lesioned Primate Model. PLOS ONE, 10(7), e0132064. 10.1371/journal.pone.0132064

Tremblay, L., Hollerman, J. R., & Schultz, W. (1998). Modifications of Reward Expectation-Related Neuronal Activity During Learning in Primate Striatum. Journal of Neurophysiology, 80(2), 964–977. 10.1152/jn.1998.80.2.964

Tremblay, L., & Schultz, W. (1999). Relative reward preference in primate orbitofrontal cortex. Nature, 398(6729), 704–708. 10.1038/19525

Van Hoesen, G. W., Pandya, D. N., & Butters, N. (1975). Some connections of the entorhinal (area 28) and perirhinal (area 35) cortices of the rhesus monkey. II. Frontal lobe afferents. Brain Research, 95(1), 25–38. 10.1016/0006-8993(75)90205-X

Wallis, J. D. (2007). Orbitofrontal Cortex and Its Contribution to Decision-Making. Annual Review of Neuroscience, 30(1), 31–56. 10.1146/annurev.neuro.30.051606.094334

Wallis, J. D., & Miller, E. K. (2003). Neuronal activity in primate dorsolateral and orbital prefrontal cortex during performance of a reward preference task. European Journal of Neuroscience, 18(7), 2069–2081. 10.1046/j.1460-9568.2003.02922.x

White, J. K., Bromberg-Martin, E. S., Heilbronner, S. R., Zhang, K., Pai, J., Haber, S. N., & Monosov, I. E. (2019). A neural network for information seeking. Nature Communications, 10(1), 5168. 10.1038/s41467-019-13135-z

Wilson, R. C., Takahashi, Y. K., Schoenbaum, G., & Niv, Y. (2014). Orbitofrontal Cortex as a Cognitive Map of Task Space. Neuron, 81(2), 267–279. 10.1016/j.neuron.2013.11.005

Xiang, J.-Z., & Brown, M. W. (1998). Differential neuronal encoding of novelty, familiarity and recency in regions of the anterior temporal lobe. Neuropharmacology, 37(4–5), 657–676. 10.1016/S0028-3908(98)00030-6

Xie, Y., Nie, C., & Yang, T. (2018). Covert shift of attention modulates the value encoding in the orbitofrontal cortex. eLife, 7, e31507. 10.7554/eLife.31507

Xu, H. A., Modirshanechi, A., Lehmann, M. P., Gerstner, W., & Herzog, M. H. (2021). Novelty is not surprise: Human exploratory and adaptive behavior in sequential decision-making. PLOS Computational Biology, 17(6), e1009070. 10.1371/journal.pcbi.1009070

Xu, N., Harnett, M. T., Williams, S. R., Huber, D., O’Connor, D. H., Svoboda, K., & Magee, J. C. (2012). Nonlinear dendritic integration of sensory and motor input during an active sensing task. Nature, 492(7428), 247–251. 10.1038/nature11601

Yamamoto, S., Monosov, I. E., Yasuda, M., & Hikosaka, O. (2012). What and Where Information in the Caudate Tail Guides Saccades to Visual Objects. The Journal of Neuroscience, 32(32), 11005–11016. 10.1523/JNEUROSCI.0828-12.2012

Young, L. T., Bagby, R. M., Cooke, R. G., Parker, J. D. A., Levitt, A. J., & Joffe, R. T. (1995). A comparison of Tridimensional Personality Questionnaire dimensions in bipolar disorder and unipolar depression. Psychiatry Research, 58(2), 139–143. 10.1016/0165-1781(95)02684-O

Zhang, K., Bromberg-Martin, E. S., Sogukpinar, F., Kocher, K., & Monosov, I. E. (2022). Surprise and recency in novelty detection in the primate brain. Current Biology, 32(10), 2160–2173.e6. 10.1016/j.cub.2022.03.064

Zhu, X. O., & Brown, M. W. (1995). Changes in neuronal activity related to the repetition and relative familiarity of visual stimuli in rhinal and adjacent cortex of the anaesthetised rat. Brain Research, 689(1), 101–110. 10.1016/0006-8993(95)00550-A

Zhu, X. O., Brown, M. W., & Aggleton, J. P. (1995). Neuronal Signalling of Information Important to Visual Recognition Memory in Rat Rhinal and Neighbouring Cortices. European Journal of Neuroscience, 7(4), 753–765. 10.1111/j.1460-9568.1995.tb00679.x

