## Supplementary Figures for "Temporal–orbitofrontal pathway regulates choices across physical reward and visual novelty"

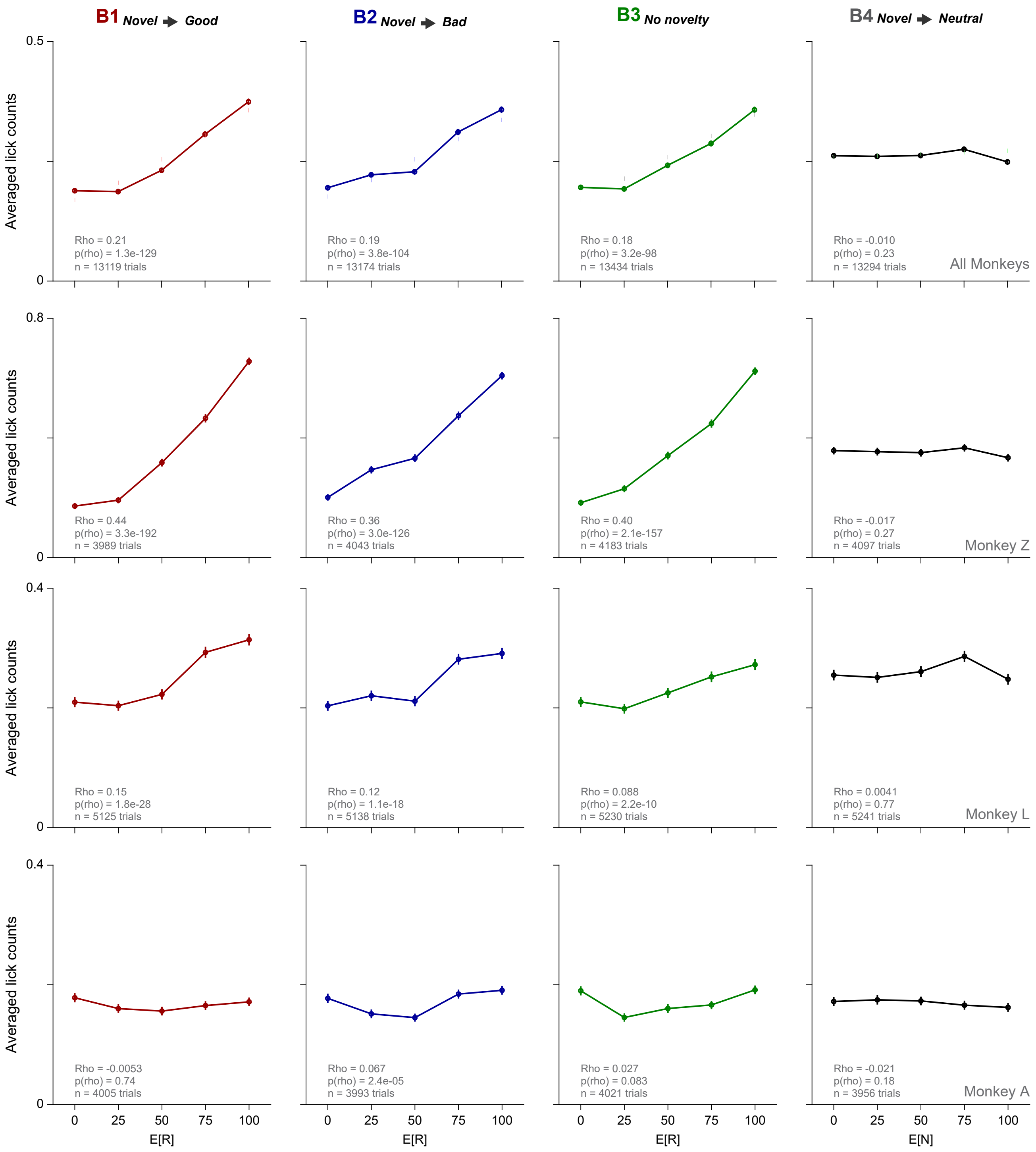


Supplementary Figure 1

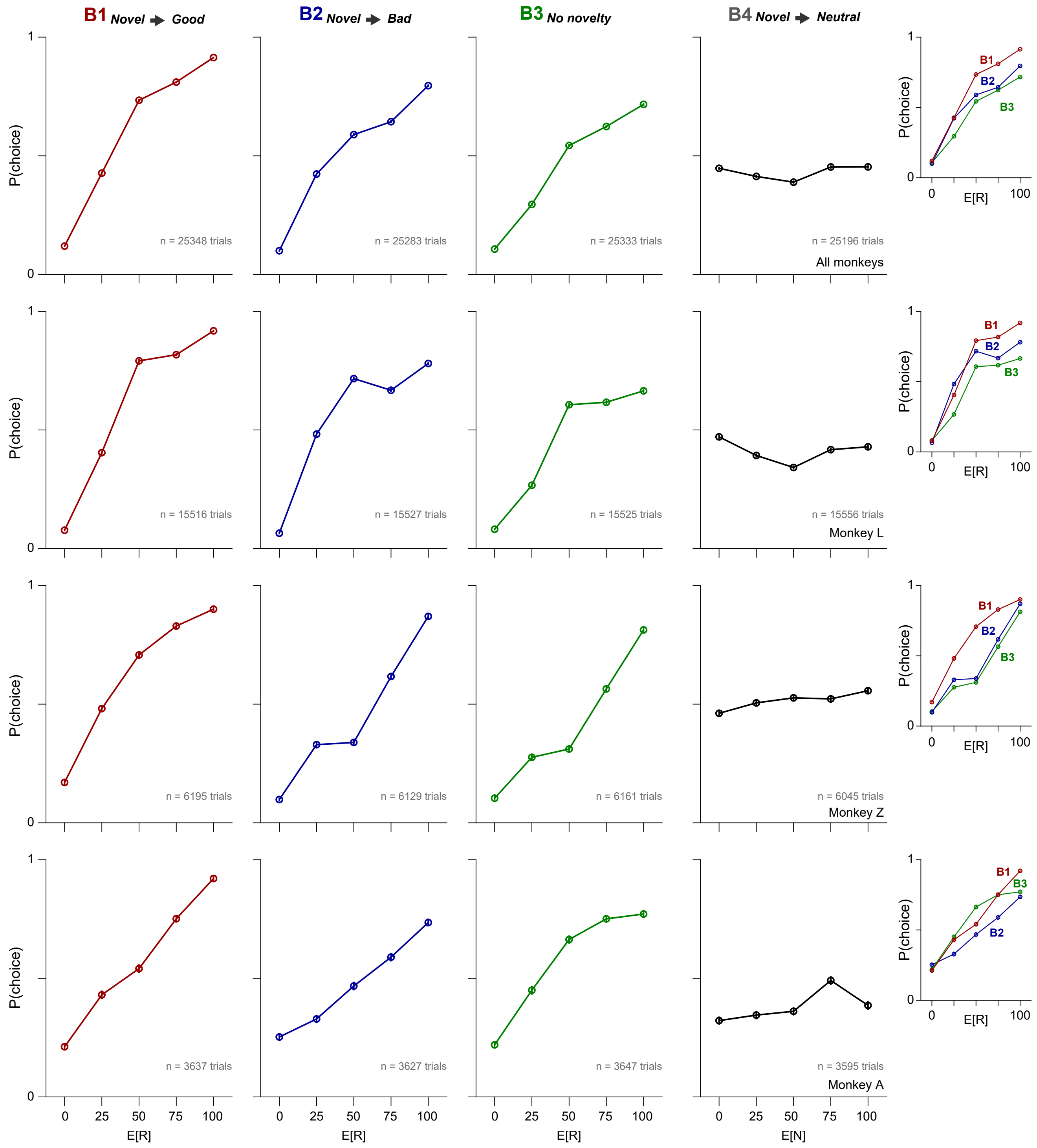


Supplementary Figure 2

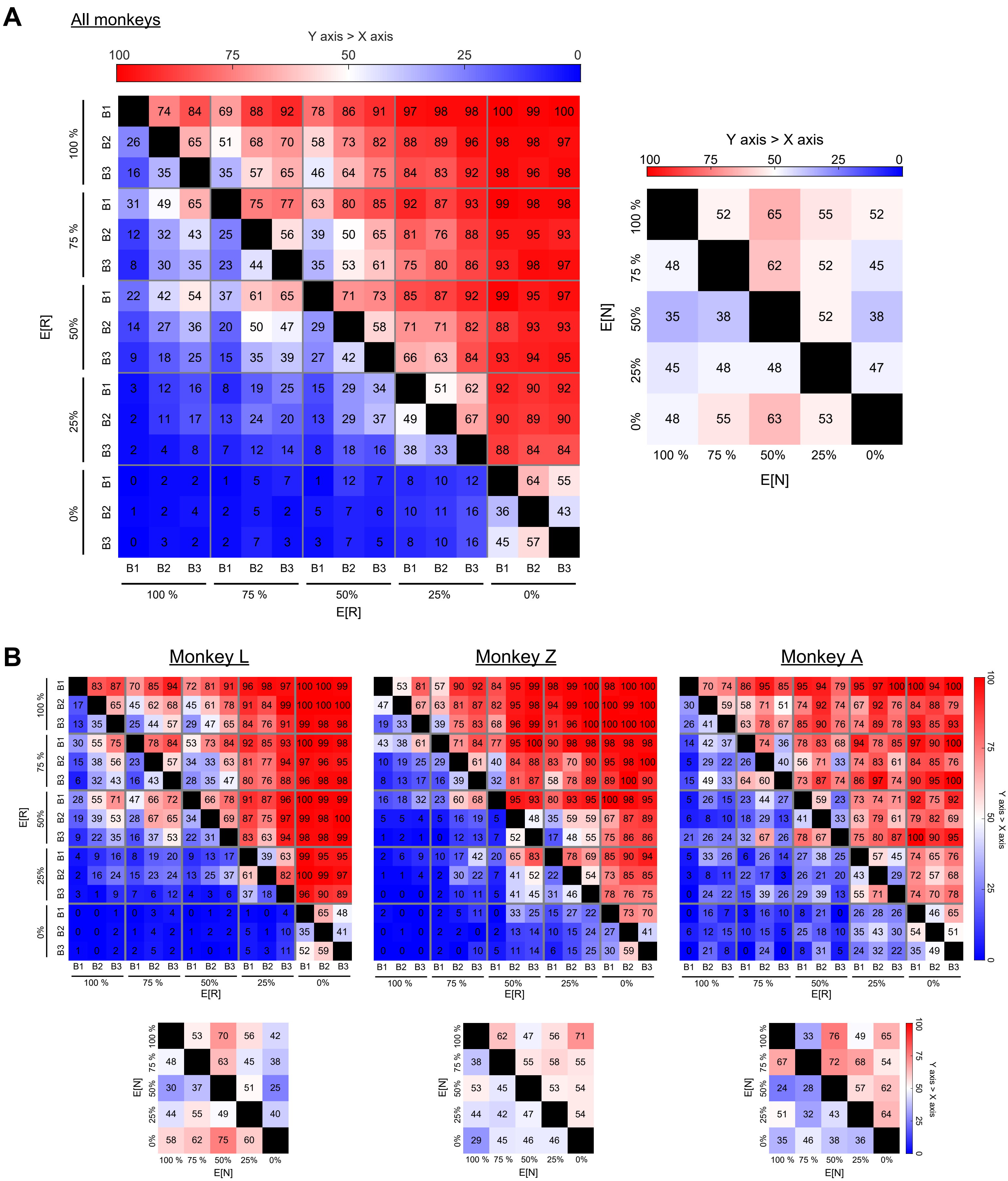


Supplementary Figure 3

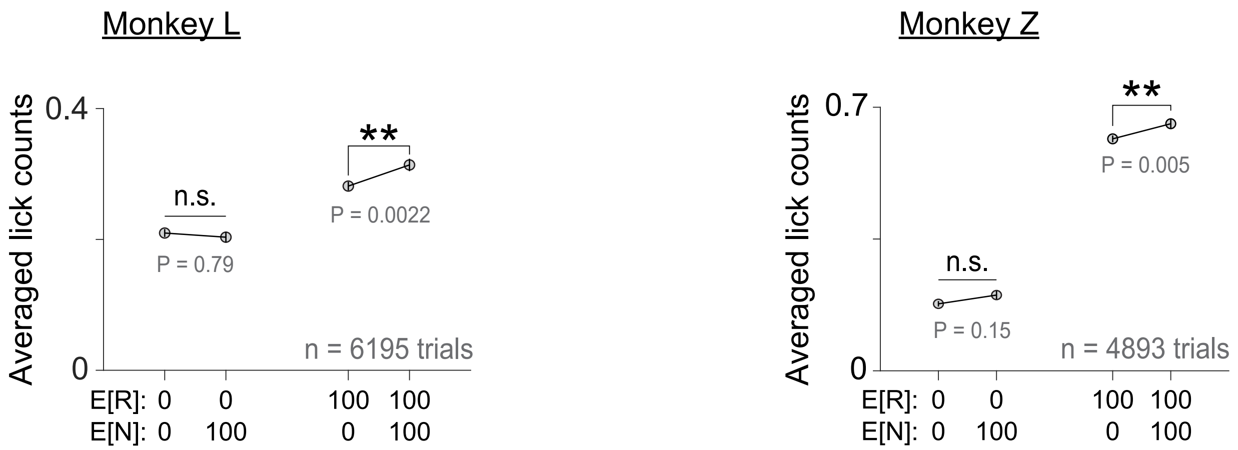


Supplementary Figure 4

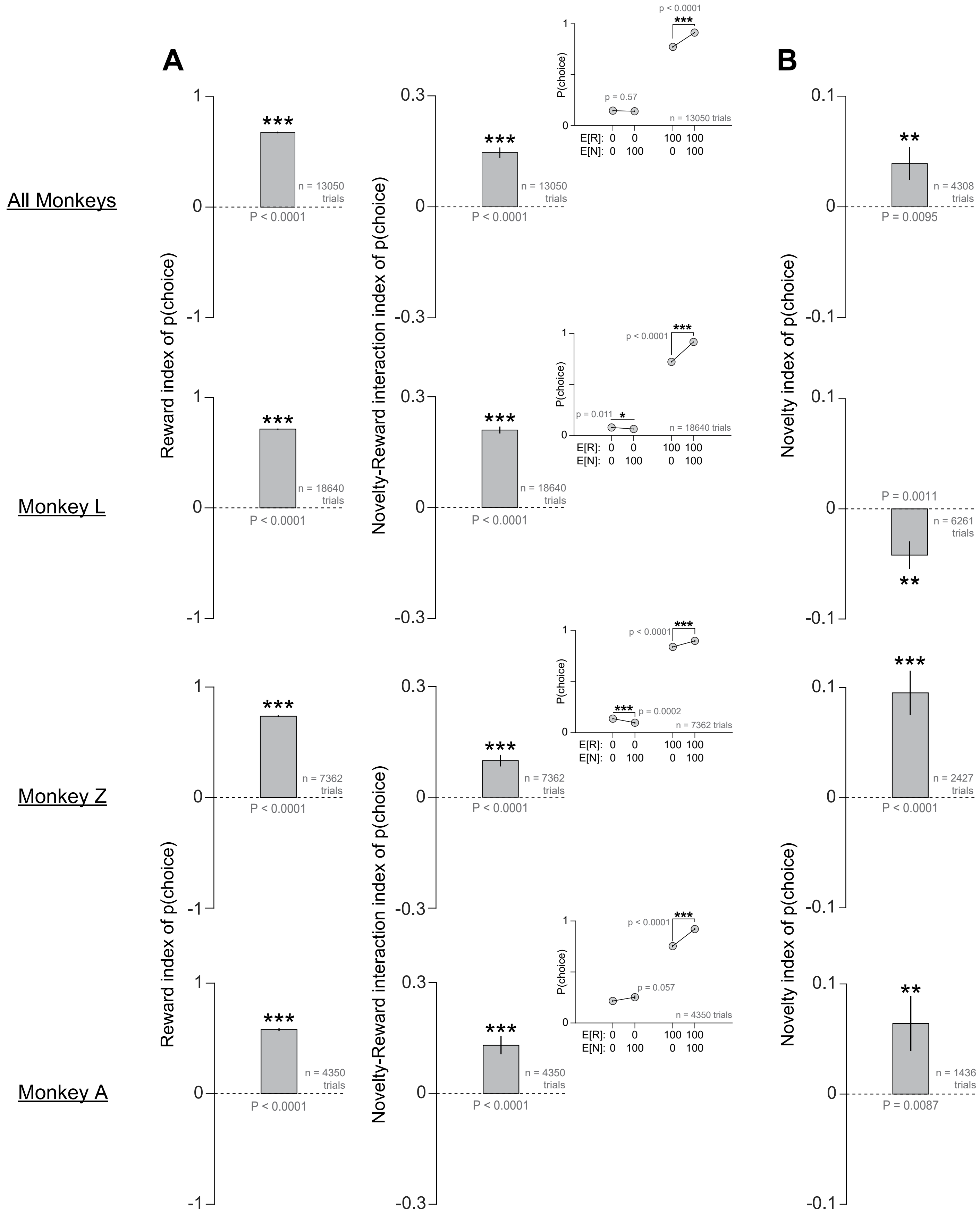


Supplementary Figure 5

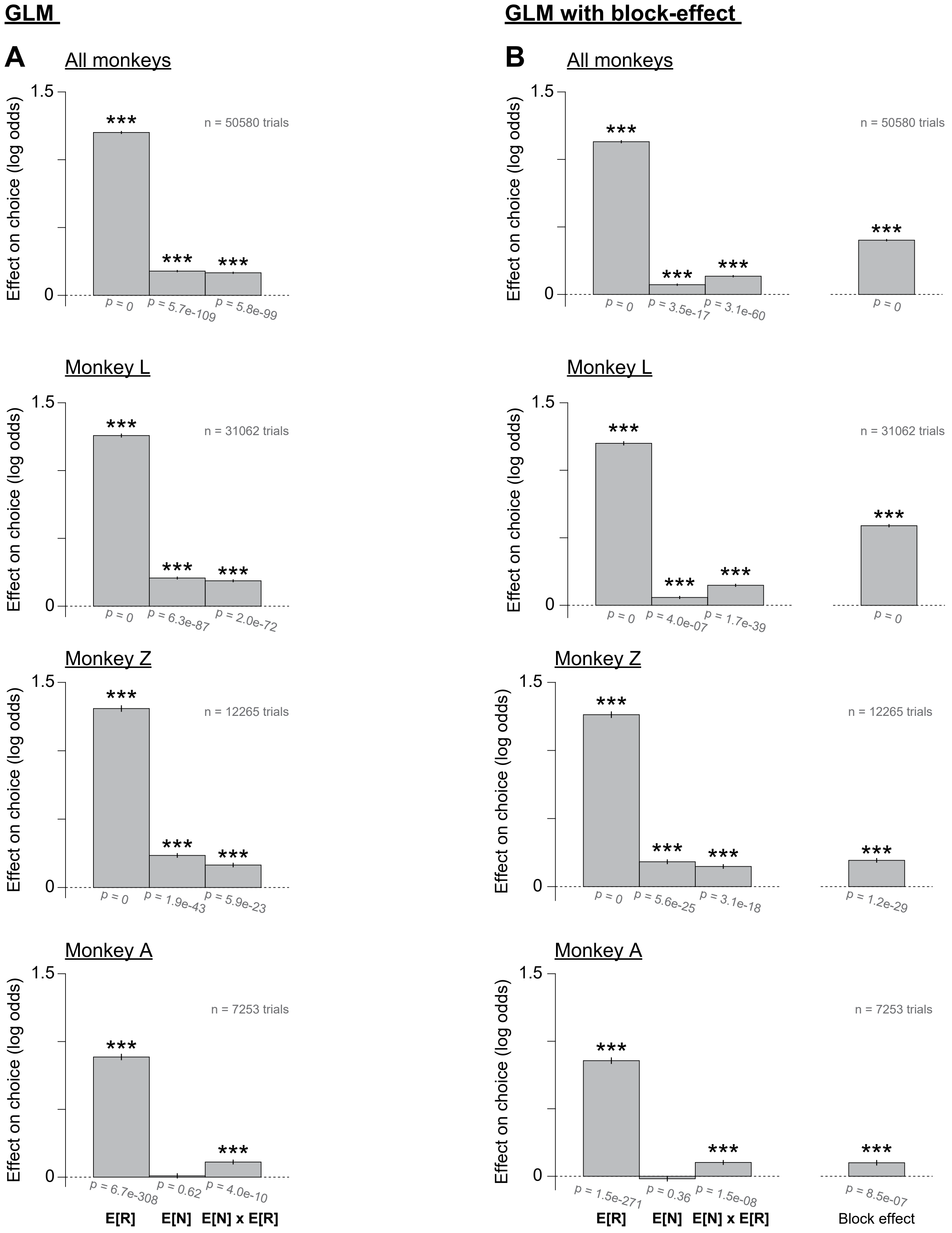


Supplementary Figure 6

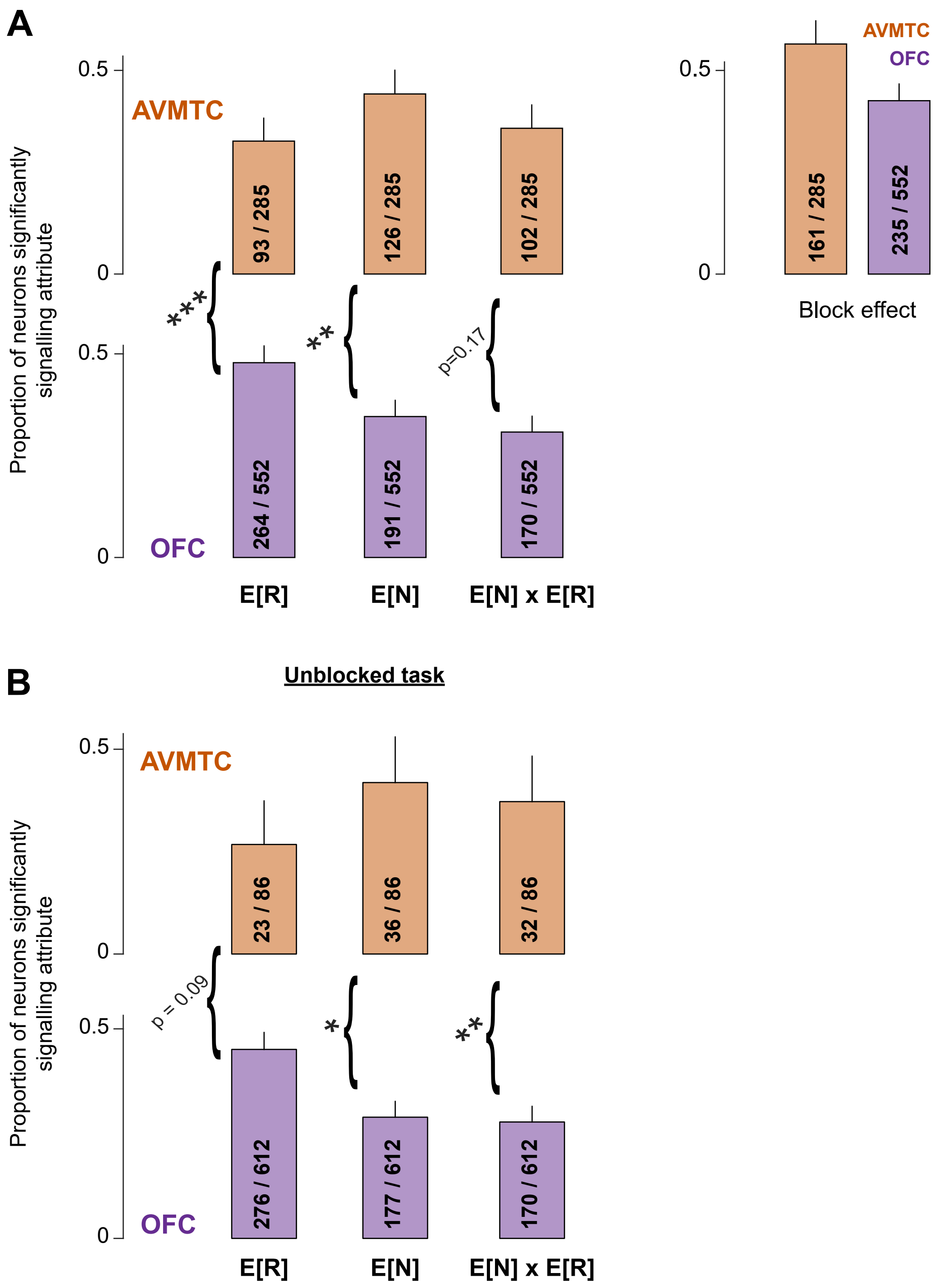


Supplementary Figure 7

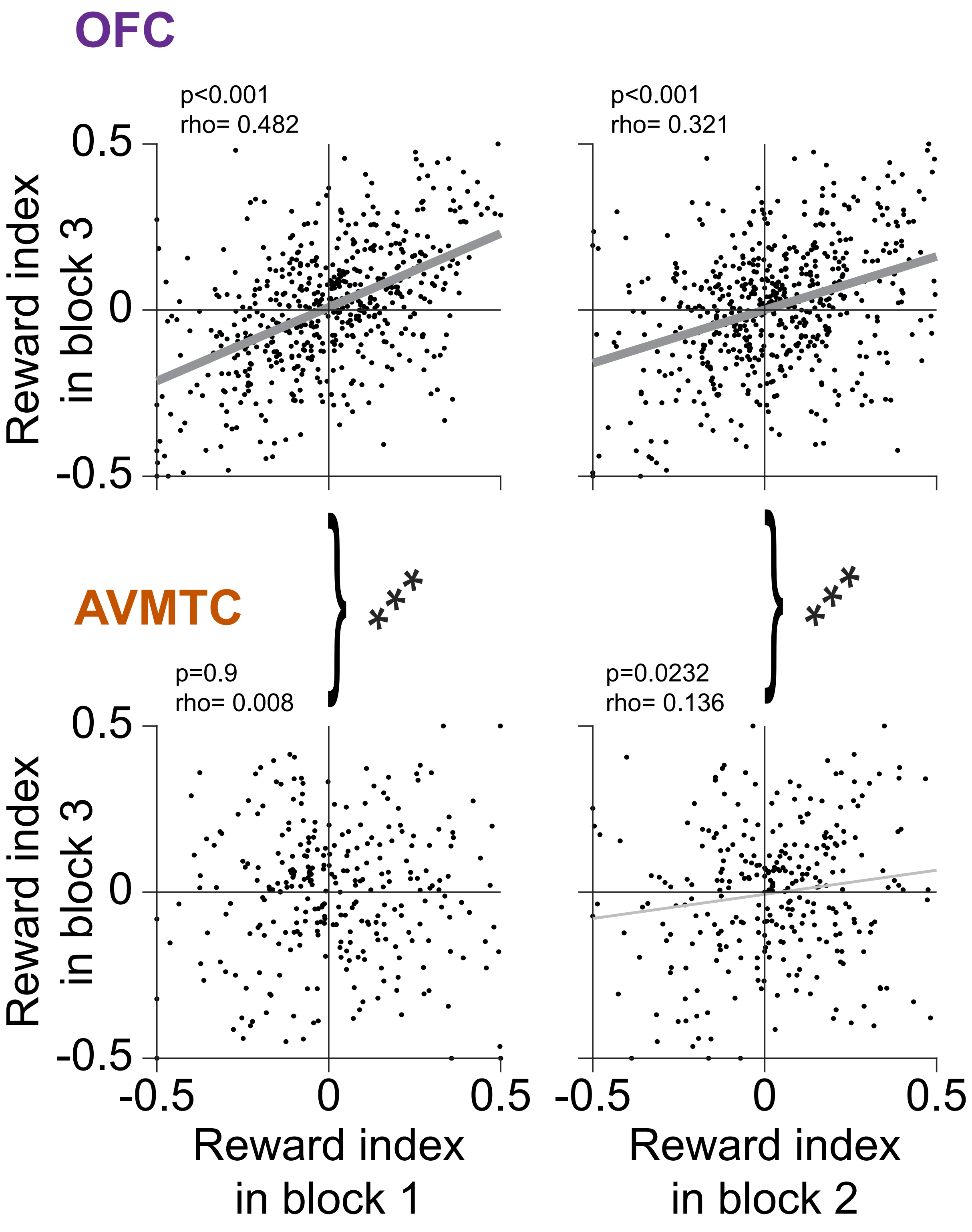


Supplementary Figure 8

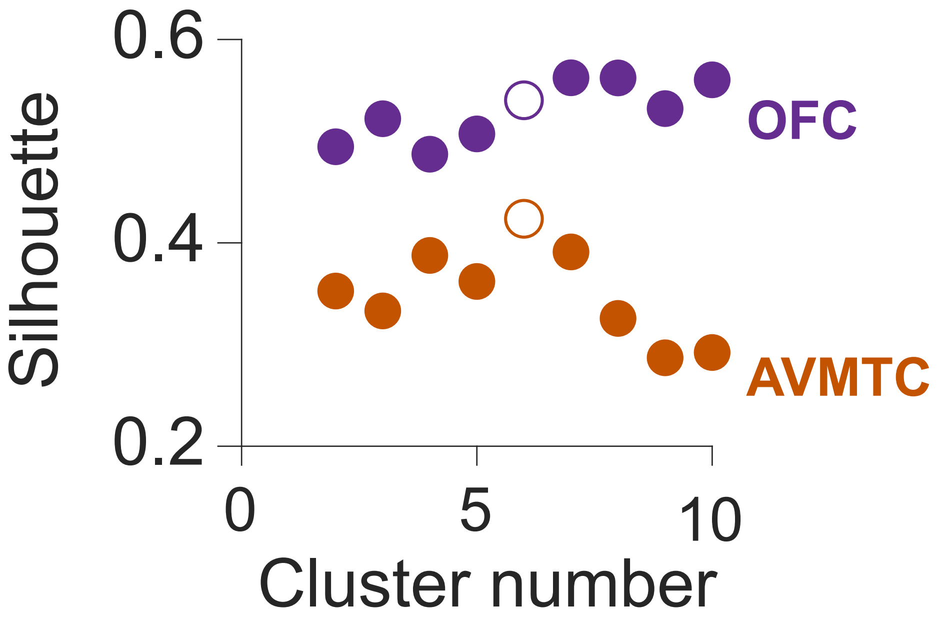


Supplementary Figure 9

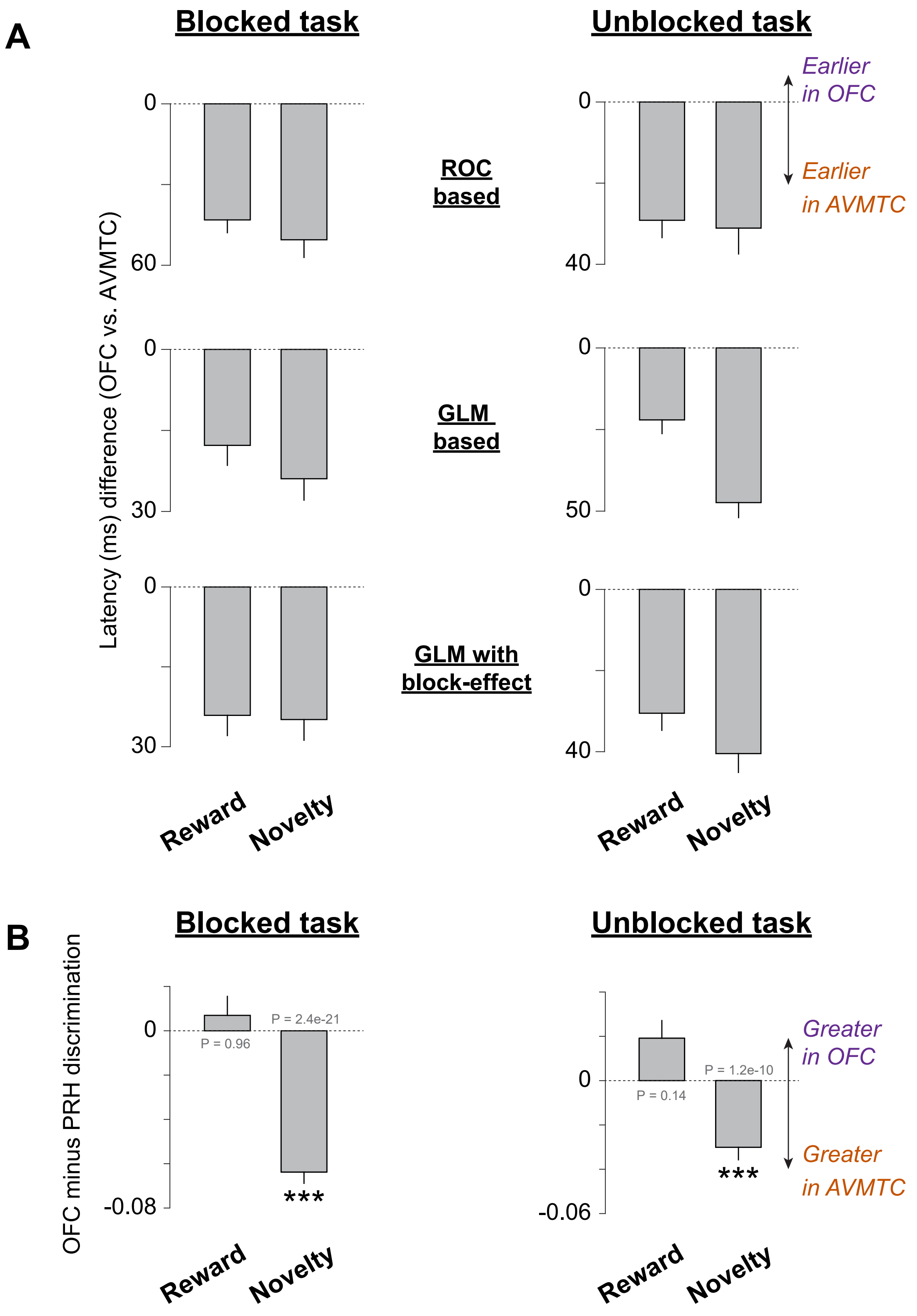


Supplementary Figure 10

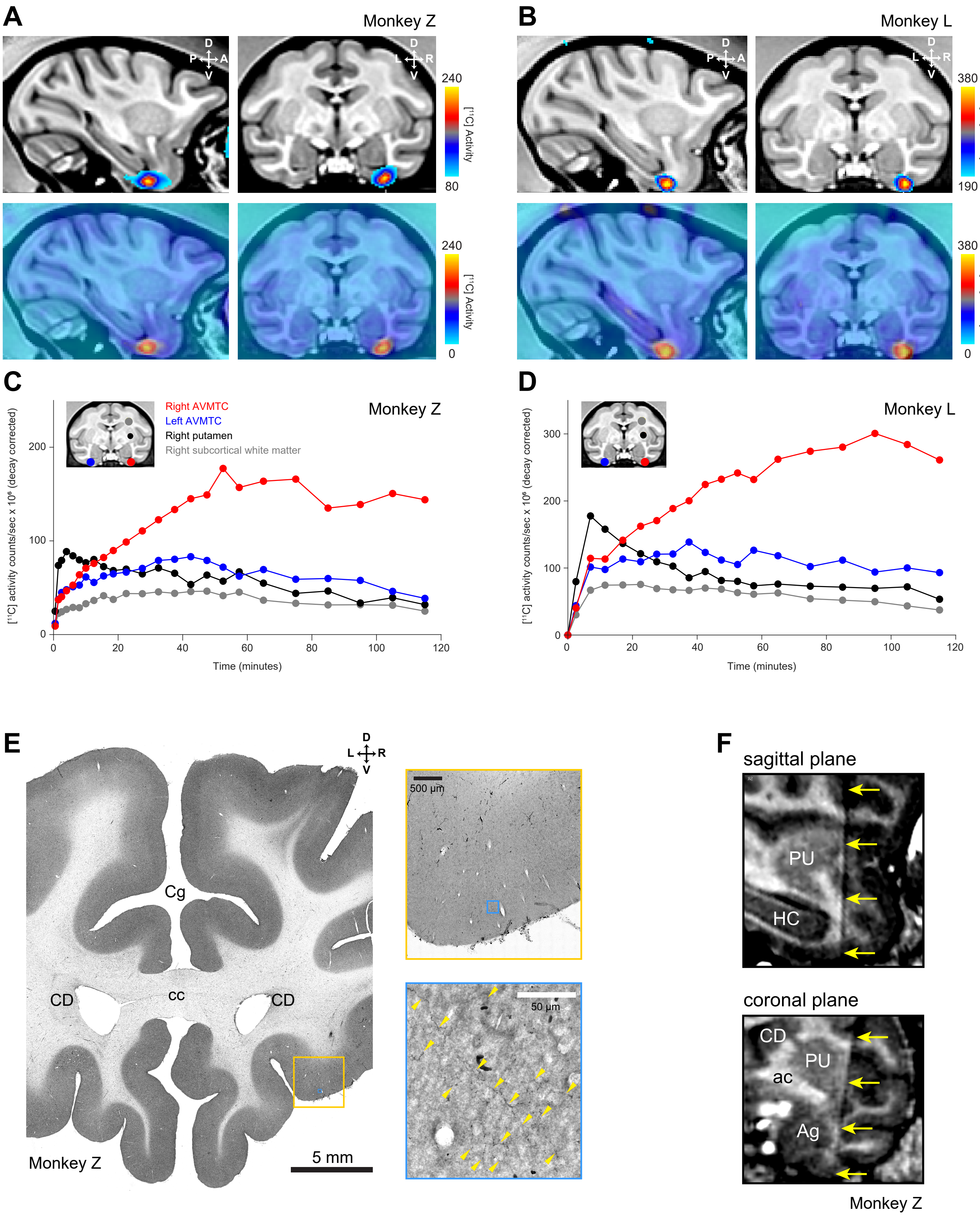


Supplementary Figure 11

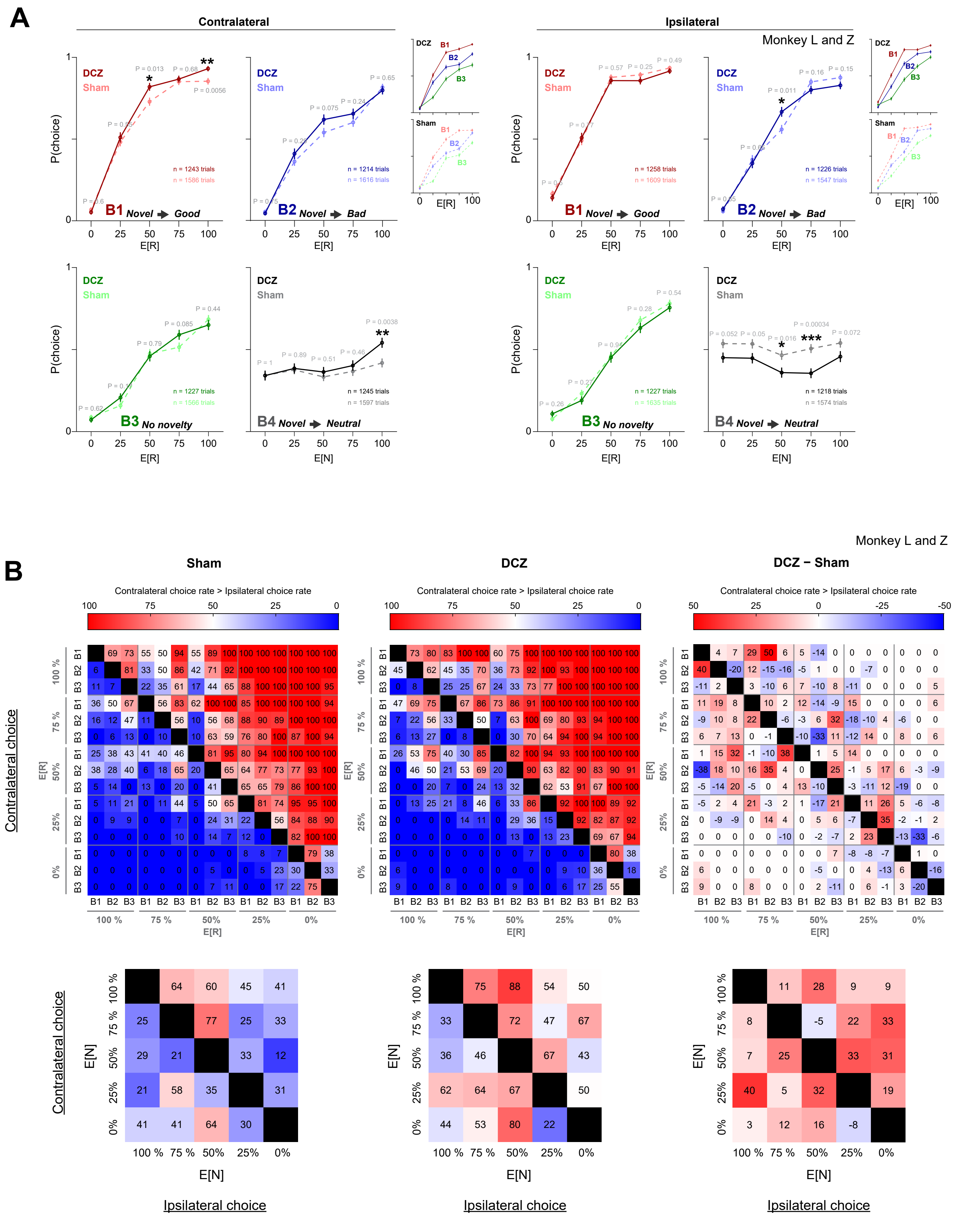


Supplementary Figure 12

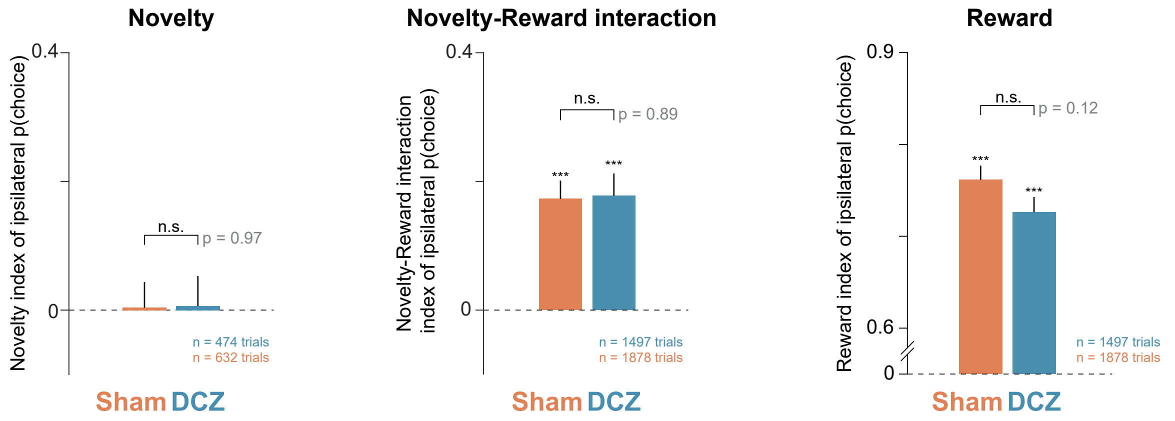


Supplementary Figure 13

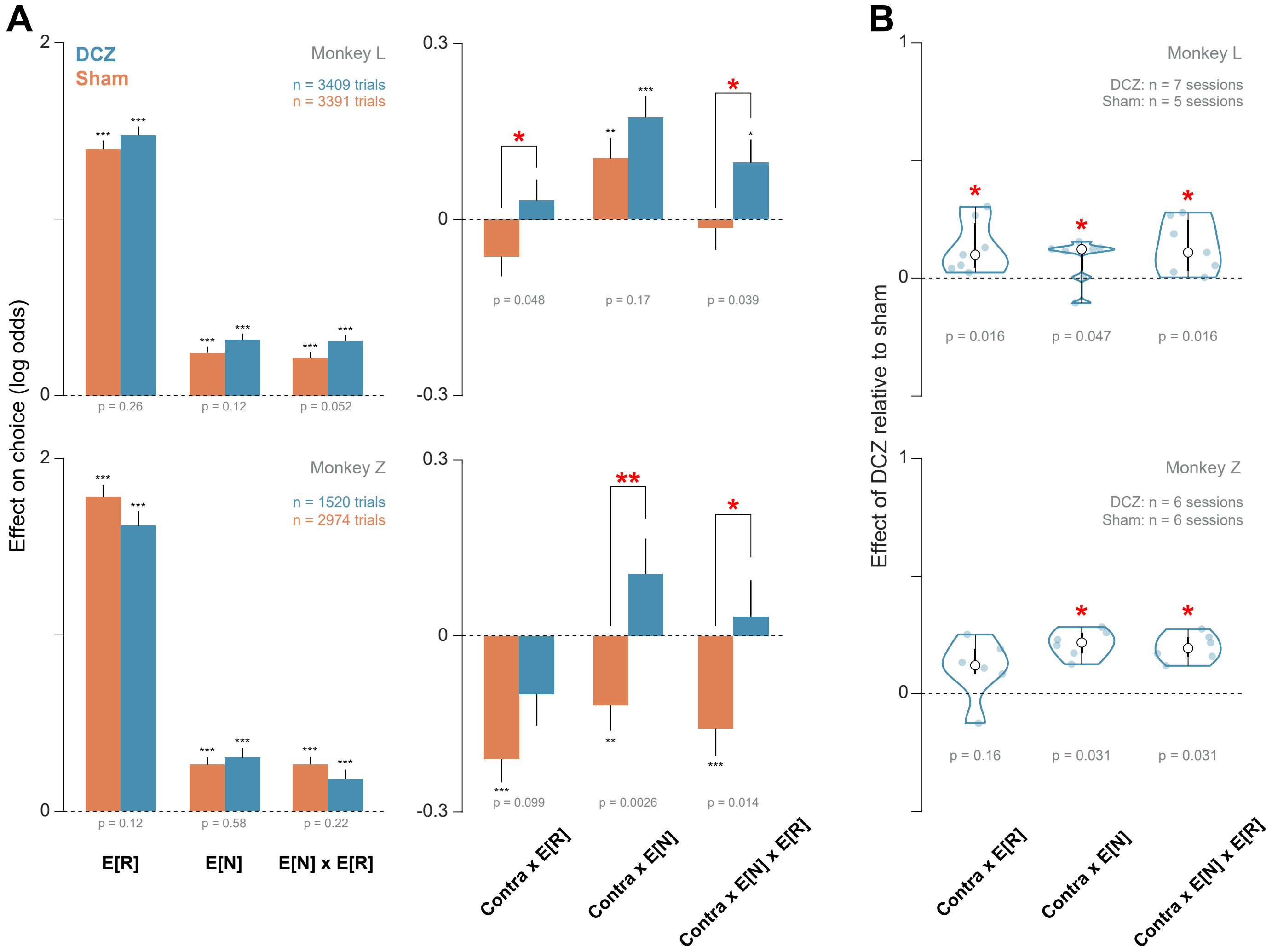


Supplementary Figure 14

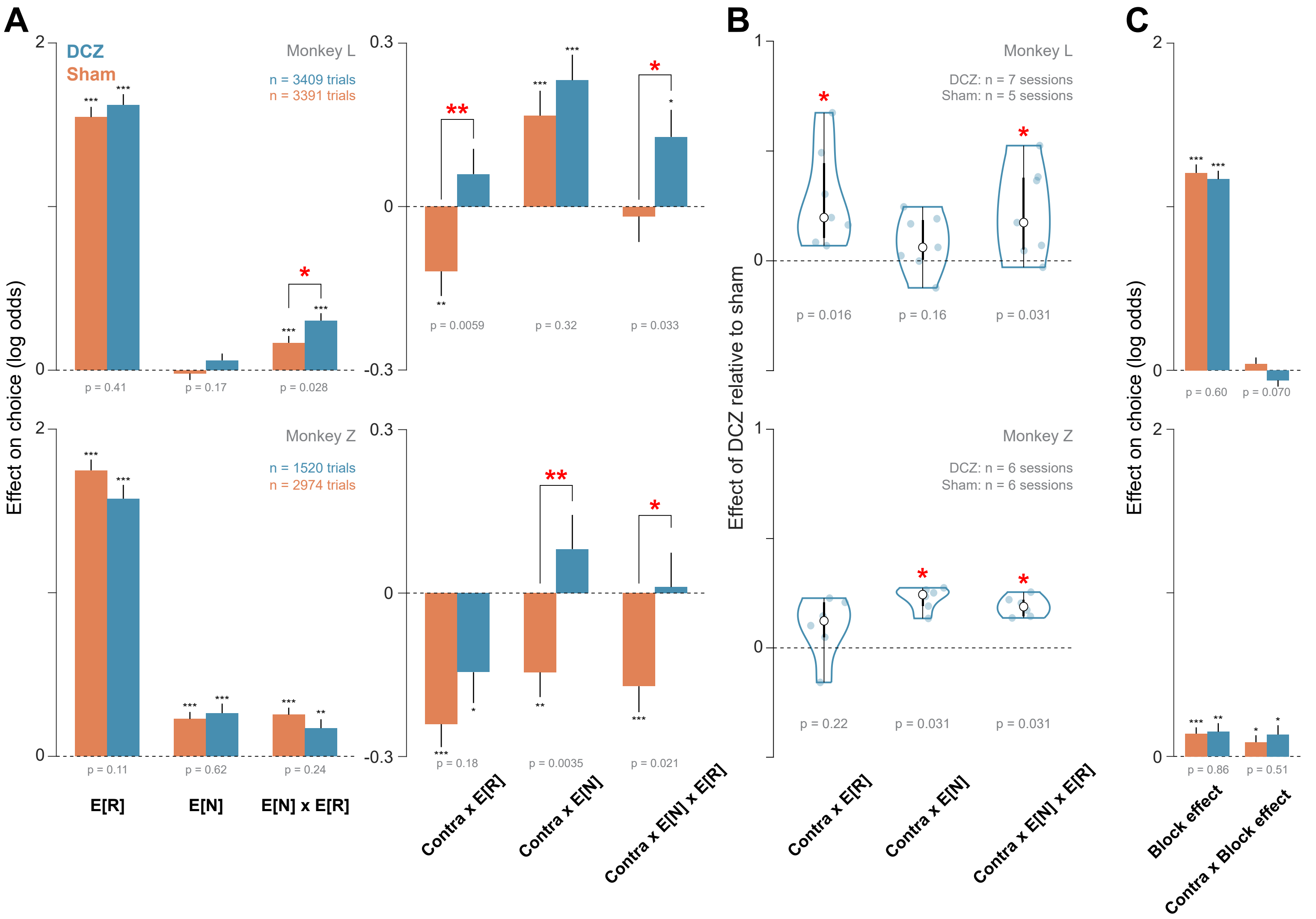


Supplementary Figure 15

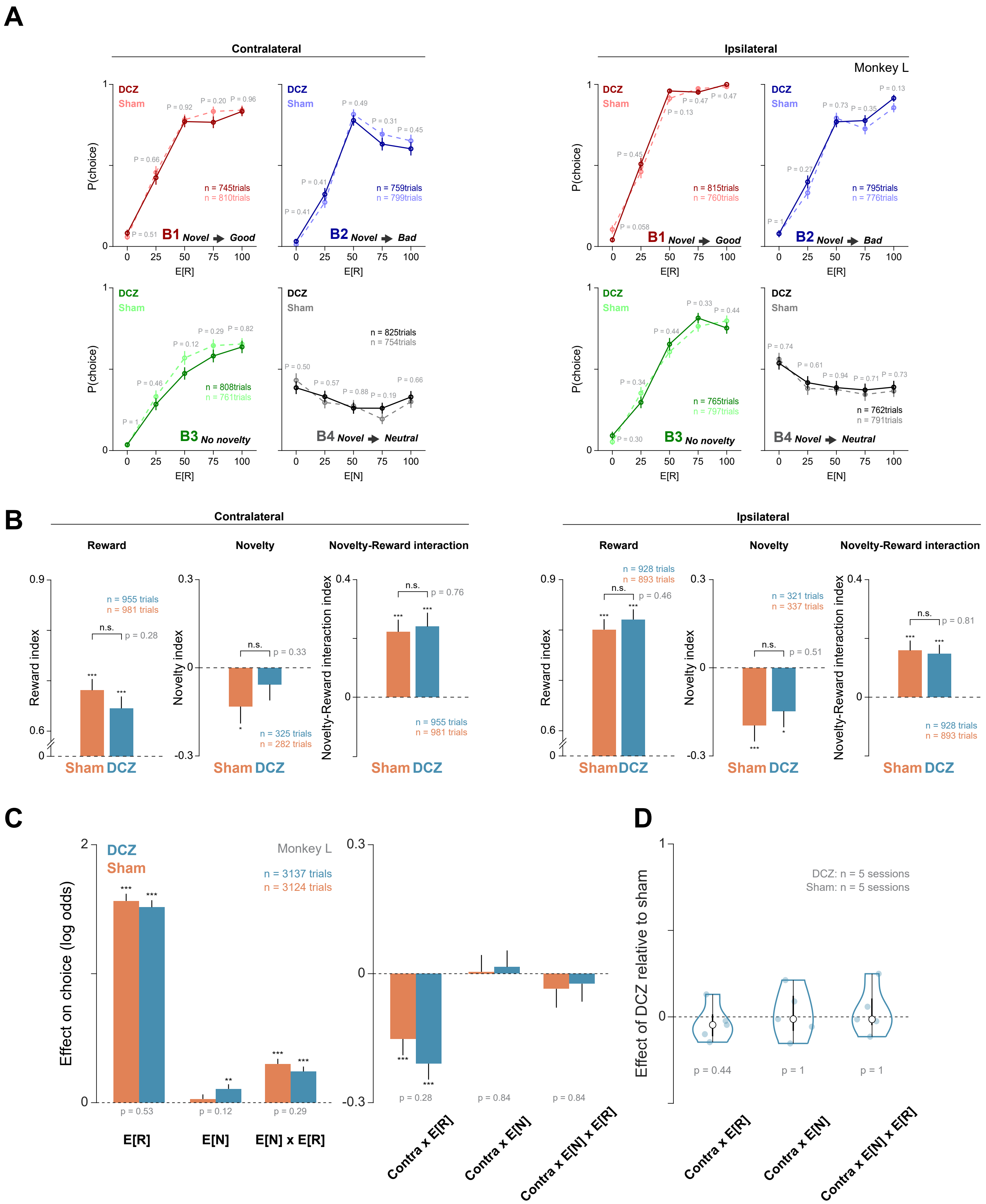


Supplementary Figure 16
